# Dispersal limitation and fire feedbacks maintain mesic savannas in Madagascar

**DOI:** 10.1101/2020.01.14.905208

**Authors:** Nikunj Goel, Erik Van Vleck, Julie C. Aleman, A. Carla Staver

## Abstract

Madagascar is regarded by some as one of the most degraded landscapes on Earth, with estimates suggesting that 90% of forests have been lost to indigenous Tavy farming. However, the extent of this degradation has been challenged: paleoecological data, phylogeographic analysis, and species diversity maps indicate that pyrogenic savannas in Central Madagascar pre-date human arrival, even though rainfall is sufficient to allow forest expansion into Central Madagascar. These observations raise a question—if savannas in Madagascar are not anthropogenic, how then are they maintained in regions where the climate can support forest? Observation reveals that the savanna-forest boundary coincides with a dispersal barrier—the escarpment of the Central Plateau. Using a stepping-stone model, we show that in a limited dispersal landscape, a stable savanna-forest boundary can form due to fire-vegetation feedbacks. This novel phenomenon, referred to as range pinning, could explain why eastern lowland forests have not expanded into the mesic savannas of the Central Highlands. This work challenges the view that highland savannas in Madagascar are derived by human-lit fires and, more importantly, suggests that partial dispersal barriers and strong non-linear feedbacks can pin biogeographical boundaries over a wide range of environmental conditions, providing a temporary buffer against climate change.

## Introduction

Savannas—defined by co-dominance of grasses and trees—cover 70% of Africa (Menaut 1983), spanning over a wide range of climatic conditions. Despite their ubiquity, mesic savannas are widely regarded as a climatic anomaly because they overlap with forest over a wide range of rainfall (1000 to ~ 2000 mm MAP) (Hirota et al. 2011, Lehmann et al. 2011, Staver et al. 2011b). The mechanism by which this climatic overlap is maintained has been a long-standing debate in biome ecology (Bond et al. 2003, Bond et al. 2005, Staver et al. 2011a) with direct implications for our conceptual understanding of which ecological processes determine tropical plant distributions and how biomes will respond to ongoing global change. Madagascar provides a unique vantage point for examining this debate, potentially offering novel theoretical insights into the spatial maintenance of global tropical biomes.

Historically, biogeographers argued that biomes occur in well-defined climatic envelopes, such that the spatial limits of biomes were thought to be solely determined by climate (Holdridge 1947, Woodward 1987). Based on this view, early naturalists in Madagascar argued that because forests are widespread in eastern lowlands, climatically similar regions in Central Highlands must have been once covered by forest, concluding that overlap in the climatic ranges of savanna and forest biomes was a result of Tavy farming (de La Bâthie 1921, Humbert 1927). Some biogeographers even argued that before human arrival, ~10.6 kyrs BP (Hansford et al. 2018), 90% of Madagascar was covered with forest (Humbert 1927)—much more than the 9% observed today (Table 1). Under this view, pyrogenic savannas in Central Madagascar were considered to be degraded ecosystems, derived from human-lit fires. Based on this reasoning, the state of Madagascar passed anti-fire policies to curb fire-setting by farmers and pastoralists (Kull 2000, 2004), and many international conservation organizations formulated plans to ‘reforest’ the Central Highlands (WRI 2014).

**Table 1:** Forest cover and deforestation estimates in Madagascar. TLA = total land area of Madagascar.

|  | Historical estimates from<br>Klein (2002) | Estimates based on satellite<br>data from Mayaux et al. (2000) |
| --- | --- | --- |
| **Pre-human forest extent (*I*)** | 90% TLA | 21% TLA |
| **Forest extent in 2000 AD** | 9% TLA | 9% TLA |
| **Total forest loss (*II*)** | 81% TLA | 12% TLA |
| **Deforestation $\left(\frac{II}{I} \times 100\right)$** | 90% | 57% |

However, an alternative viewpoint—based on three independent lines of empirical evidence—indicates pyrogenic savannas in Central Madagascar pre-date human arrival. First, charcoal sediments and *C*_4_ pollen in paleo-cores show frequent fire activity in Central Highlands [in Fig. 1A, we collate paleo sites from published sources; also see Table 1 and Metadata S1]. In fact, the oldest record of savanna vegetation from Lake Tritrivakely in Central Madagascar precedes human arrivals in Madagascar by 6.4 kyrs BP (Gasse and Van Campo 1998). Second, grass and forb endemism in Madagascar is much higher than the global average for large islands (Bond et al. 2008, Vorontsova et al. 2016); moreover, faunal collections reveal the coevolution of many open habitat specialists with the surrounding pyrogenic vegetation (Bond et al. 2008). Accumulation of such diverse biota is considered unlikely if mesic savannas have a recent anthropogenic origin. Third, phylogeographic analysis show ancient genetic divergence (~55 kyrs BP) between lemur populations dwelling in the eastern and western sides of the island (Yoder et al. 2016). Since trees are critical for lemur dispersal to maintain gene flow, the ancient divergence has been interpreted as evidence for the absence of a continuous forest habitat before human arrival. Together, these lines of evidence suggest that pyrogenic savannas in Madagascar pre-date human arrival and that forest cover in Madagascar before human arrival was much less than 90% of the total land area. Current estimates based on satellite imagery (Green and Sussman 1990, Mayaux et al. 2000) and historical forest maps (Humbert and Cours Darne 1965, Koechlin 1972, Bond et al. 2008) put early 20^*th*^ century forest cover around 21% of Madagascar (see Table 1 and Fig. 1A), implying that the observed overlap in the climatic ranges of mesic savannas and forests may not be an anthropogenic artifact.

**Figure 1:**
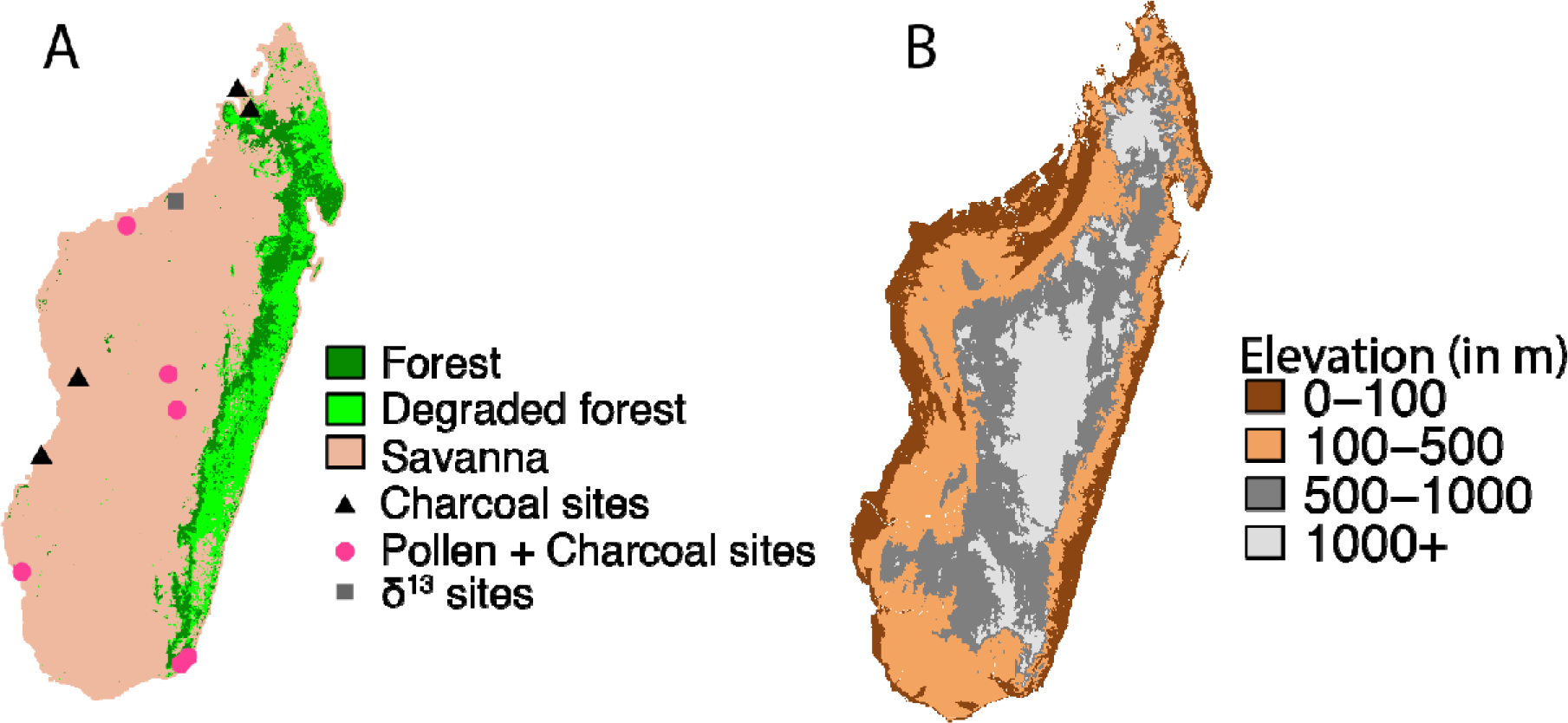
Remotely sensed distribution of savanna and forest biomes in Madagascar (Mayaux et al. 2000) (A), and an elevation map of Madagascar (Jarvis et al. 2008) (B). In plot (A), dots indicate the locations of paleo sites in Madagascar (see Table S1 and Appendix S1 for more details on paleoecological data). Note that the eastern edge of the Central Plateau is coincident with the savanna-forest boundary. We hypothesize that the Central Plateau could have prevented the expansion of forest into Central Madagascar due to dispersal limitation.

These empirical findings present a theoretical challenge: if the overlap in the climatic ranges of biomes pre-dates human arrival, how are wet savannas maintained in Central Madagascar, given that these regions are climatically suited to allow forest expansion? Clearly, bioclimatic (one-climate-one-biome) models are insufficient in explaining savanna distribution in Madagascar. A developing school of thought is that savanna and forest are alternative stable states maintained by positive fire-vegetation feedbacks (Staver et al. 2011a). Based on this view, savannas in the Central Highlands could be considered as alternative biome state to forest (Staver et al. 2011b). However, recent work using one-dimensional diffusion models suggests that in a landscape with a precipitation gradient, the bistability disappears, and, like bioclimatic models, the biome boundary is solely determined by climate (Wuyts et al. 2017). Diffusion models, however, too, have their own limitations: diffusion approximations fail when density-dependent feedbacks are strong and dispersal is limited (Keitt et al. 2001), as is the case for biomes in Madagascar, where the savanna-forest boundary is coincident with a dispersal barrier—the eastern edge of the Central Plateau (Fig. 1B). However, only a handful of theoretical (Goldberg and Lande 2007) and empirical studies (see Rapoport 1982, Gaston 2003) examine the role of barriers on the propagation of populations.

Invasion theory with Allee effect may offer useful insights on how tropical biome boundaries interact with climate and dispersal barriers, since the mathematical structure and analysis of these invasion models (Murray 2001) is very similar to the spatial models of savanna and its alternative biome state, forest (Goel et al. 2018, Wuyts et al. 2019). Using Ulbrich’s data on the dispersal of Muskrat, Rapoport (1982) noted that the species expanded at a constant rate in a homogeneous landscape, as predicted by the diffusion models, but the invasion briefly paused when it encountered a topographical barrier. Invasion theory suggests these time lags may result from Allee effect (Taylor and Hastings 2005)—when dispersal from the preceding patch is slow, the population density in the focal patch may take a while to cross the Allee threshold, but once the threshold is crossed, the population jumps to the carrying capacity. Using a stepping stone invasion model (the discrete analog of diffusion model), Keitt et al. (2001) and follow up work by Wang et al. (2019) showed that when positive feedbacks are strong and dispersal is weak, the lag time approaches infinity. In other words, the ecotone stabilizes even though the climate could still permit expansion. This phenomenon, referred to as ‘range pinning’ in invasion theory, provides a plausible theoretical mechanism by which tropical forests in Madagascar might be restricted to the eastern lowland.

Using a stepping stone model that captures climate-dependent fire feedbacks with variation in landscape porosity, we analyze the role of dispersal barriers in determining the spatial distribution of savanna-forest biomes. In particular, *(i)* we ask whether the escarpment of the Central Plateau could have maintained mesic savannas in Central Highlands by preventing forest expansion via dispersal limitation. To evaluate the robustness of our theoretical results, *(ii)* we also simulate biome distributions using present-day rainfall (Huffman and Bolvin 2013) [Tropical Rainfall Measuring Mission (TRMM)] and topography data (Jarvis et al. 2008) [Shuttle Radar Topography Mission (SRTM)], with and without dispersal limitation.

## Modeling Framework

We model the spatial dynamics of savanna and forest in Madagascar using a stepping-stone model. In this modeling approach, space is represented as a collection of discrete patches that are arranged on a lattice. Within each patch, the vegetation dynamics are governed by local climate and fire-vegetation feedbacks. We assume adjacent patches interact via passive dispersal of seeds. Below, we first describe the local vegetation dynamics, and then the seed dispersal term.

### Local climate-fire-vegetation feedbacks via the mean-field approach

In savanna and forest ecosystems, fire-vegetation feedbacks play an important role in determining the local tree cover in landscape (Trapnell 1959, Swaine et al. 1992). When tree cover is low, fire spreads readily (Archibald et al. 2009, Staver and Levin 2012) and perpetually maintains the landscape in low tree cover (Higgins et al. 2000, Moreira 2000, Prior et al. 2010, Veenendaal et al. 2018). However, if tree cover increases past some threshold, the grass matrix becomes discontinuous, and fire ceases to spread (Hennenberg et al. 2006, Archibald et al. 2009, Pueyo et al. 2010). Consequently, trees form a closed canopy and exclude the remaining grass layer by shading it out (Hennenberg et al. 2006, Lloyd et al. 2008). Here, we model these fire-vegetation interactions using a mean-field approach within a patch.

For mathematical simplicity, we assume the landscape consists of only forest trees and grasses (Goel et al. 2018). In doing so, we ignore savanna trees, such that results provide intuition for the boundary between savanna/grasslands and forest, rather than tree cover within these ecosystems. We also assume that grasses colonize on much faster time scales than trees (Purata 1986). This assumption has three consequences: (*i*) the bare ground can be ignored since grasses quickly colonize any patch not occupied by a competitively dominant forest tree; (*ii*) since the landscape consists of only forest trees and grasses, the sum of the local density of forest trees (*T*) and grasses (*G*) can be set to a constant (see Table S2 for parameter definitions), here taken to be unity (*i.e.*, *T* + *G* = 1); (*iii*) the instantaneous dynamics of biomes can be expressed in terms of tree density alone, such that the instantaneous local grass cover is given by *G* = 1 − *T*.

Dynamically, we assume trees establish at a rate proportional to the product of local tree density (to approximate local seed production) and local grass density (the empty spaces for colonization), with precipitation *P* as the proportionality constant (see also Staver et al. 2011a, Staver and Levin 2012, Touboul et al. 2018). This proportionality constant could potentially be modified to incorporate edaphic constraints on tree establishment, although this extension is not shown here. However, results would not change qualitatively, as soils only act to change the strength of fire-feedbacks (see Bowman and Perry 2017). Next, we assume that the per-capita mortality rate of forest trees in response to fire is a step function *ϕ*(*T*) (we provide the functional form of *ϕ* in Fig. 2 caption), which depends on the local density of grasses and, thus, trees (*G=1−T*; see also Staver et al. 2011a, Staver and Levin 2012). The mortality rate *ϕ* is high at high grass density (such as in an open-canopy savanna/grassland, where fire spreads readily) and low at low grass density (such as in a closed-canopy forest). Combining these demographic processes yields a mean-field growth function of trees:

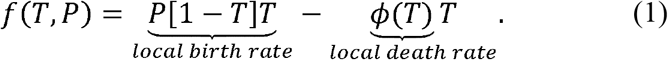

**Figure 2:**
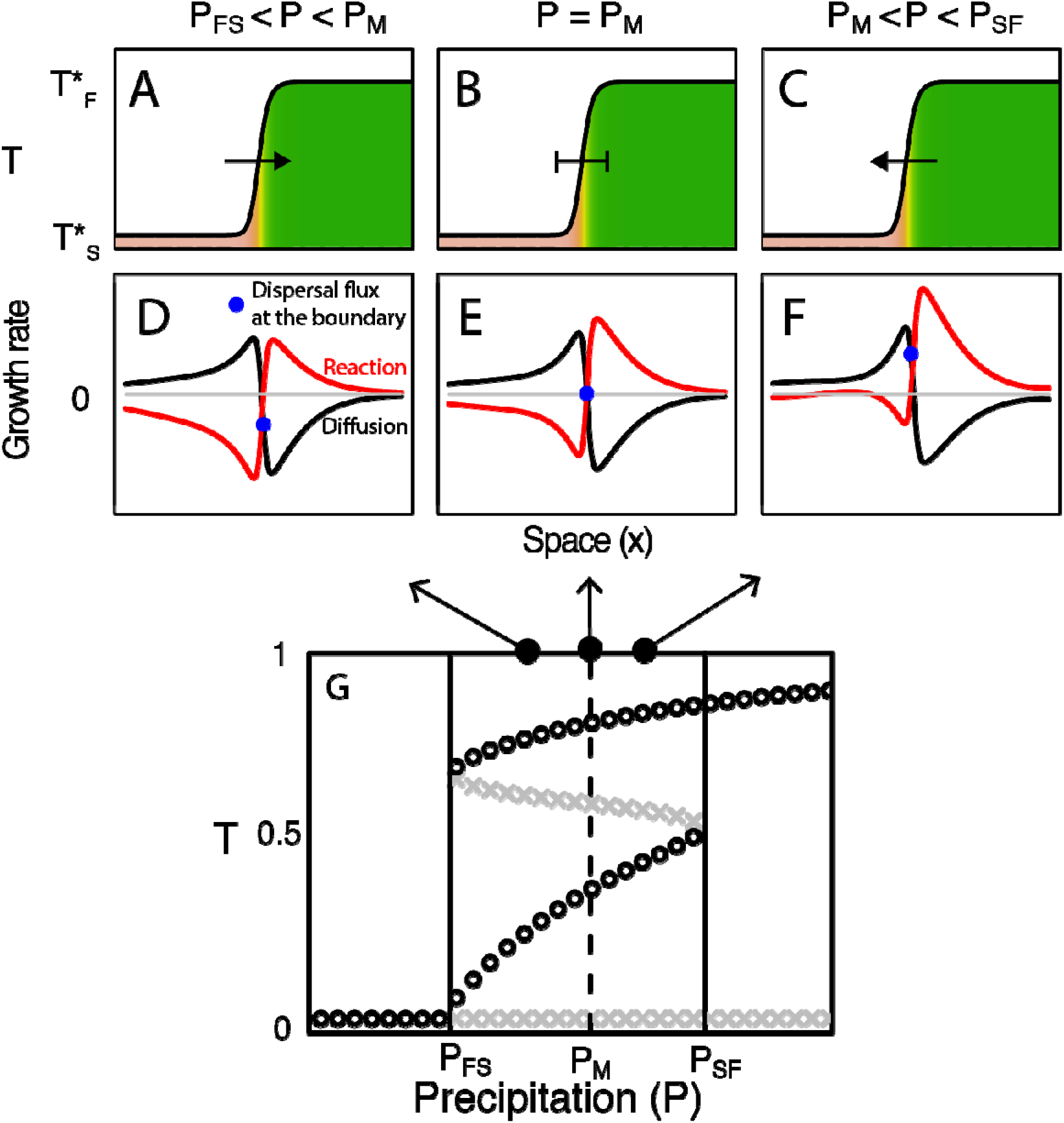
Direction of boundary movement (A-C) as a function of dispersal flux (D-F) for different precipitation values in the bistable region (G). In the bistable region(*P*_*FS*_ < *P* < *P*_*SF*_), the reaction term has two stable roots—savanna and forest (black circles)—which are separated by an unstable root (grey crosses). In the second row, the black and red lines are the reaction and diffusion terms in Eq. 3, and the blue circle represents the dispersal flux at the boundary. In the 1D diffusion model, when *P*_*FS*_ < *P* < *P*_*M*_,savanna invades forest due to negative dispersal flux at the boundary (A and D). Conversely, when *P*_*M*_ < *P* < *P*_*SF*_, forest invades savanna due to positive dispersal flux (C and F). A stationary savanna-forest boundary is formed only when dispersal flux is exactly zero (E). This condition is met at a unique precipitation value—Maxwell Precipitation *P*_*M*_ (B). Thus, in a 1D landscape with precipitation gradient, the savanna-forest boundary equilibrates at *P*_*M*_. We use 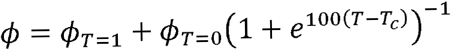 to generate the bifurcation diagram, where ϕ_*T*=1_ = 0.13,ϕ_*T*=0_ = 0.23, and *T*_c_ = 0.55 (threshold tree cover for fire spread). Plots D, E, and F were generated for *P* = 0.48, 0.53(*P*_*M*_), and 0.58, respectively. Simulations in D-F were performed on a one-dimensional lattice of size 500 with *D* = 50. For the separameter values, we get *P*_*FS*_ = 0.377,*P*_*M*_ = 0.53, and *P*_*SF*_ = 0.668.

Phenomenologically, this mean-field model captures two important aspects of tropical biomes. First, in the intermediate precipitation range, *P*_*FS*_ < *P* < *P*_*SF*_, fire-vegetation feedbacks are substantial. In this precipitation interval, the *f*(*T*,*P*) has two stable roots—savanna (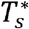) and forest (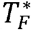) states (black circles in Fig. 2G)—and an unstable root 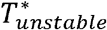 (grey crosses in Fig. 2G). This unstable root represents the threshold tree-cover below which the local grass cover becomes continuous, and fire spread readily. Second, outside the bistable region, the fire feedbacks are absent, and the system equilibrates to a climatically determined equilibrium: when *P* > *P*_*SF*_ (critical precipitation value for savanna to forest transition) forest is stable and, conversely, when *P* < *P*_*FS*_ (critical precipitation value for forest to savanna transition) savanna is stable.

### Dispersal via stepping-stone approach

We assume that the dispersal (number of seeds dispersed per unit time) between adjacent patches is proportional to the density of trees in each patch, such that the proportionality rate constant is a function of the patch itself. Mathematically, we express this assumption using a stepping-stone approach in which patches *i* and *i* + 1 exchange seeds at a rate proportional to *D*_*i*_. Intuitively, *D*_*i*_ is the net influx of seeds in patch *i* when the tree cover differs in the focal and succeeding patch by unity (henceforth referred to as dispersal rate). Similarly, patches *i* − 1 and *i* exchange seeds at a rate proportional to *D*_*i*−1_. Thus, in a 1D landscape, the net dispersal flux at patch *i* from both directions is Φ_*i*_ = *D*_*i*_[*T*_*i*+1_ − *T*_*i*_] + *D*_*i*−1_[*T*_*i*−1_ − *T*_*i*_] (Humphries et al. 2011). We assume the topographical features of the landscape determine *D*_*i*_: when the landscape is flat *D*_*i*_ takes a high value and decreases as the surface inclines. In a 2D landscape, Φ_*i*,*j*_ = *D*_*i*,*j*_[*T*_*i*+1,*j*_ − 2*T*_*i*,*j*_ + *T*_*i*,*j*+1_] + *D*_*i*−1,*j*_[*T*_*i*−1,*j*_ − *T*_*i*,*j*_ + *D*_*i*,*j*−1_[*T*_*i*,*j*−1_ − *T*_*j*_] (Hoffman et al. 2017). This 2D the dispersal flux term is the sum of two 1D dispersal flux terms corresponding to each spatial coordinate.

Combining the local vegetation dynamics in Eq. 1 with patch specific dispersive interactions yields a 1D heterogeneous stepping-stone model:

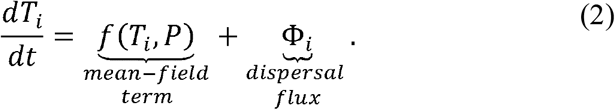

To obtain a 2D version of the model, we replace the subscript [*i*] by [*i*,*j*], representing *x* and *y* coordinates. Stepping-stone models in Eq. 2 are widely used ecology to describe spatial dynamics such as range limits (Keitt et al. 2001) and animal movement (Okubo 1986). Past theoretical work suggests that Eq. 2 has a traveling-wave solution (Keitt et al. 2001, Wang et al. 2019), in which the spatial profile of the tree cover resembles a chain of falling dominos (Keitt et al. 2001, Humphries et al. 2011). In this analogy, each domino represents a vegetation patch: standing and fallen dominoes represent forest and savanna patches, respectively, and the cascading front represents the savanna-forest boundary.

## Results

In the following sections, we consider two versions of the stepping-stone model: a homogeneous dispersal model with a constant dispersal rate, and a heterogeneous dispersal model in which dispersal rate is a function of the slope of the terrain. For each model, we (*i*) present a graphical approach to describe ecological conditions that yield a stable boundary and (*ii*) simulate biome patterns in Madagascar using a parameterized model that incorporates information on both present-day rainfall and topography.

Unfortunately, neither version of the stepping-stone model can be solved analytically for mean-field growth function in Eq. 1. Therefore, we consider a piecewise linear growth function (McKean 1970) in the Appendix S1 to analytically explore the effects of dispersal barriers on the position of the boundary as a function of precipitation and dispersal constant (Fig. S1, S2, and S3). Although the piecewise linear function is not biologically motivated, it captures the bistability of savanna and forest biomes, which is central to explaining the main results of this paper and offers a more comprehensive insight that can be tested by simulating realistic functional form of fire-vegetation feedbacks, *e.g.*, Eq. 1. We refer our readers to Humphries et al. (2011) and Marsden et al. (1993) for a detailed discussion on the analytical properties of a stepping-stone model presented in the Appendix S1.

### Homogeneous dispersal model

#### Theory

We begin with a uniform dispersal model with isotropic dispersal rate *D* that does not depend on the topographical features of the landscape. This simplification allows us to express the net dispersal flux (Φ_*i*_) as *D*[*T*_*i*+1_ − 2*T*_*i*_ + *T*_*i*−1_], which, in the high-*D* limit, can be further simplified to a diffusion approximation to yield a 1D reaction-diffusion model:

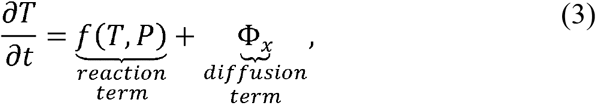

where Φ_*x*_ = *D*∂^2^*T*/∂*x*^2^ is the 1D Laplacian operator. By extension, for a 2D landscape, Φ_*x,y*_ = *D*[∂^2^*T*/∂*x*^2^ + ∂^2^*T*/∂*y*^2^]. The results of the diffusion model are used to establish a baseline for comparison with the heterogeneous dispersal model that is presented in the next section.

Reaction-diffusion models (Eq. 3) are an idealized representation of spatial dynamics of populations, as they ignore stochasticity in demographic rates and instead model these processes as continuous functions. This simplification is based on the idea that local stochastic processes can be approximated as continuous functions at large spatial scales (Skellam 1951, Levin 1992). This methodology can provide an intuitive understanding of boundary dynamics, including tropical biome boundaries (Wuyts et al. 2017, Goel et al. 2018). Here, we briefly discuss the results of the diffusion model in the context of biomes in Madagascar (see Goel et al. 2018 for details).

According to Eq. 3, the dynamics at the boundary are governed by local fire-vegetation feedbacks (reaction term) and dispersal flux from adjacent patches (diffusion term). By defining the position of the boundary as 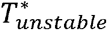, we can neglect the contribution of fire feedbacks at the boundary because at equilibrium, the reaction term is always zero, *i.e.*, 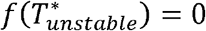. However, because this equilibrium is unstable, even a small dispersal flux Φ_*x*_ at the boundary can initiate a traveling wave of one biome state invading other. When Φ_*x*_ is negative (Fig. 2D), savanna invades forest (Fig. 2A), and, conversely, when Φ_*x*_ is positive (Fig. 2F), forest invades savanna (Fig. 2C). The boundary equilibrates only when the Φ_*x*_ is exactly zero (Fig. 2E), a condition which is met at a unique precipitation value, referred to as Maxwell Precipitation (*P*_*M*_) (Fig. 2B). Intuitively, Maxwell Precipitation is the bioclimatic limit of tropical biomes, where both savanna and forest invade each other at an equal rate. Therefore, in a landscape with a precipitation gradient, the boundary equilibrates at the spatial location that receives *P*_*M*_ (see Appendix S1 for analytical results). As mentioned in the introduction, the 1D diffusion model is analogous to bioclimatic models that predict the spatial limits of biomes are climatically determined.

Recent work shows that this 1D approximation is limited and that a more realistic representation of dispersal as a 2D process can qualitatively change the equilibrium position of the boundary (Goel et al. 2018). To a first approximation, the equilibrium position of the boundary is still determined by Maxwell precipitation contour (*P*_*MC*_; analogous to *P*_*M*_ in 2D), but it may locally deviate from *P*_*MC*_ depending on the local curvature of the *P*_*MC*_. These curvature effects arise because, in a 2D landscape, the dispersal flux is not only dependent on precipitation but also on the number of savanna and forest neighbors, which, in turn, is a function of the curvature of *P*_*MC*_. These curvature effects are a result of source-sink dynamics in which spatial subsidies from productive patches can maintain the population in sub-optimal patches.

#### Large-scale simulations

To establish a baseline prediction, we simulate vegetation in Madagascar using a calibrated homogeneous 2D diffusion model with *P*_*M*_ = 1538 mm MAP (see Appendix S1). This estimate of *P*_*M*_ was obtained from the analysis of vegetation patterns in mainland Africa (Goel et al. 2018 found P_M_ lies within 1508 ± 84 mm MAP with 1538 mm MAP as the most probable value), where the diffusion model provides more accurate distribution of biomes and is consistent with other recent empirical (Staal et al. 2016) and theoretical studies (Wuyts et al. 2017). We initialize our simulation with historical estimates of biome distributions in Madagascar in Fig. 1A (our best estimate of ‘initial conditions’ on the island). This initial condition roughly matches reconstructed biome patterns from LGM and Holocene that show forest in Madagascar were restricted to the eastern lowlands (see Buney 1996, Adams and Faure 1997a).

We find that this basic diffusion model overpredicts forest cover (Fig. 3A) and cannot explain how mesic savannas are maintained in Central Madagascar. Note that the biome boundary is not coincident with *P*_*Mc*_ because of the source-sink dynamics described above.

**Figure 3:**
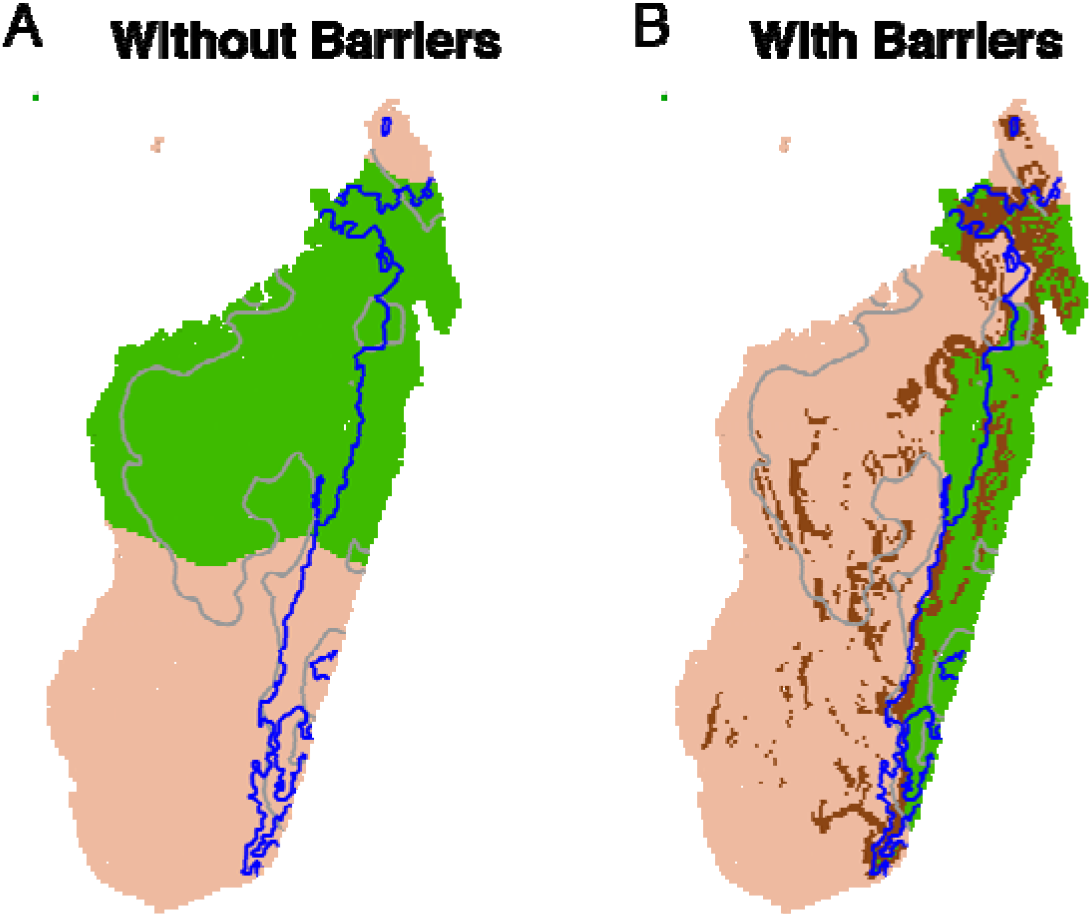
Simulated distribution of savanna and forest without (A) and with (B) dispersal barriers. The blue and grey lines indicate the forest extent from Mayaux et al. (2000) and Maxwell precipitation contour (*P*_*M*_ = 1538 mm MAP), respectively. The brown pixels in the plot (B) indicate dispersal-limited patches. The uniform dispersal model (2D diffusion model) produces extensive forest cover in Central Madagascar (A). However, when topography is included in the simulation, the savanna-forest boundary is pinned at the escarpment of the Central Plateau (B). The heterogeneous dispersal model in (B) reproduces biome patterns with 92% accuracy, which is much higher than 54% accuracy from the uniform dispersal model in (A) (see Appendix S1, Fig. S6, and Metadata S2). Thus, boundary pinning may explain how mesic savannas are ecologically maintained in CentralMadagascar.

### Heterogeneous dispersal model

#### Theory

In the above model, we assumed a homogeneous landscape with a high dispersal rate. This assumption allowed us to approximate dispersal as a continuous diffusion process. However, the diffusion approximation fails when the dispersal rate is low, as it is when the savanna-forest boundary approaches a dispersal barrier. In this section, we show that in a stepping-stone model with a low dispersal rate, a stable biome boundary can arise due to fire-vegetation feedback despite the availability of environmentally suitable patches ahead. This property of bistable traveling waves is referred to as range pinning in invasion biology (Keitt et al. 2001).

To examine how range pinning affects the boundary propagation near a dispersal barrier, we make two simplifying assumptions. First, we only consider biome dynamics in a 1D landscape. Although previous work shows that this assumption is not ideal (Goel et al. 2018), we think that it may be reasonable in Madagascar, based on the observation that both the savanna-forest boundary and the eastern slope of the Central Plateau run linearly along the eastern coast of Madagascar. This symmetrical arrangement suggests that we may be able to ignore the second dimension, but a full analytical treatment of the 2D stepping stone model can be found in Hoffman et al. (2017) (note that we also provide supporting 2D simulations after offering analytical intuition for 1D case). Second, we consider a constant dispersal rate between adjacent patches, except between patches *k* and *k* + 1, corresponding to the location of the dispersal barrier. We incorporate the barrier in our model using a binary dispersal scheme in Eq. **2**: *D*_*k*_ = *d* and *D*_*i*≠*k*_ = *D* such that *D* » *d*.

This dispersal scheme has the advantage that when the boundary is far from the barrier, the biome dynamics can be described using the diffusion approximation, *i.e.*, the equilibrium position of the boundary is uniquely determined by *P*_*M*_. However, if the boundary encounters a barrier during propagation, the diffusion approximation breaks down. As a result, the boundary may become pinned at the barrier before reaching its bioclimatic limit *P*_*M*_. To build intuition for how this mechanism could have stabilized the biome boundary at the edge of the Central Plateau, we investigate the dynamics of forest invading savanna at the dispersal barrier when *P* > *P*_*M*_.

Consider a 1D traveling wave with savanna in patches *k* and below, and forest in patches *k* + 1 and above. Since the dispersal rate at the barrier is low, the boundary savanna (patch *k*) and forest (patch *k* + 1) patches can be considered to be near their equilibrium densities. This simplification allows us to approximate dispersal flux in Eq. 2 at patch *k* as 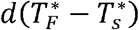. Next, using this approximation, we examine the conditions on *d* for which forest fails to invade savanna.

At patch *k*, the biome dynamics can be expressed as

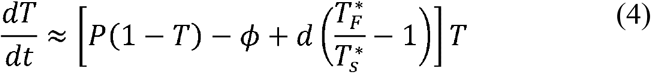

(for convenience we lose the subscript *k*). In the bistable region, the system has at most four roots (see bifurcation diagram in Fig. 2G). Since *T* = 0 is a trivial root, we direct our attention to the other three roots, which can be obtained by setting the square bracket term to zero. Graphically, finding the roots is equivalent to identifying points where growth function intersects the zero growth line. When *d* ≈ 0, 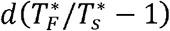 is negligible, and the dynamics at patch *k* are solely governed by the *P*(1 − *T*) − *ϕ*, which intersects the zero growth line at two stable equilibrium points—savanna and forest—and at an unstable equilibrium 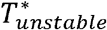 (black curve in Fig. 4A). Since the patch is initially in the savanna state, fire feedbacks maintain the patch in the savanna attractor and, consequently, forest invasion fails.

**Figure 4:**
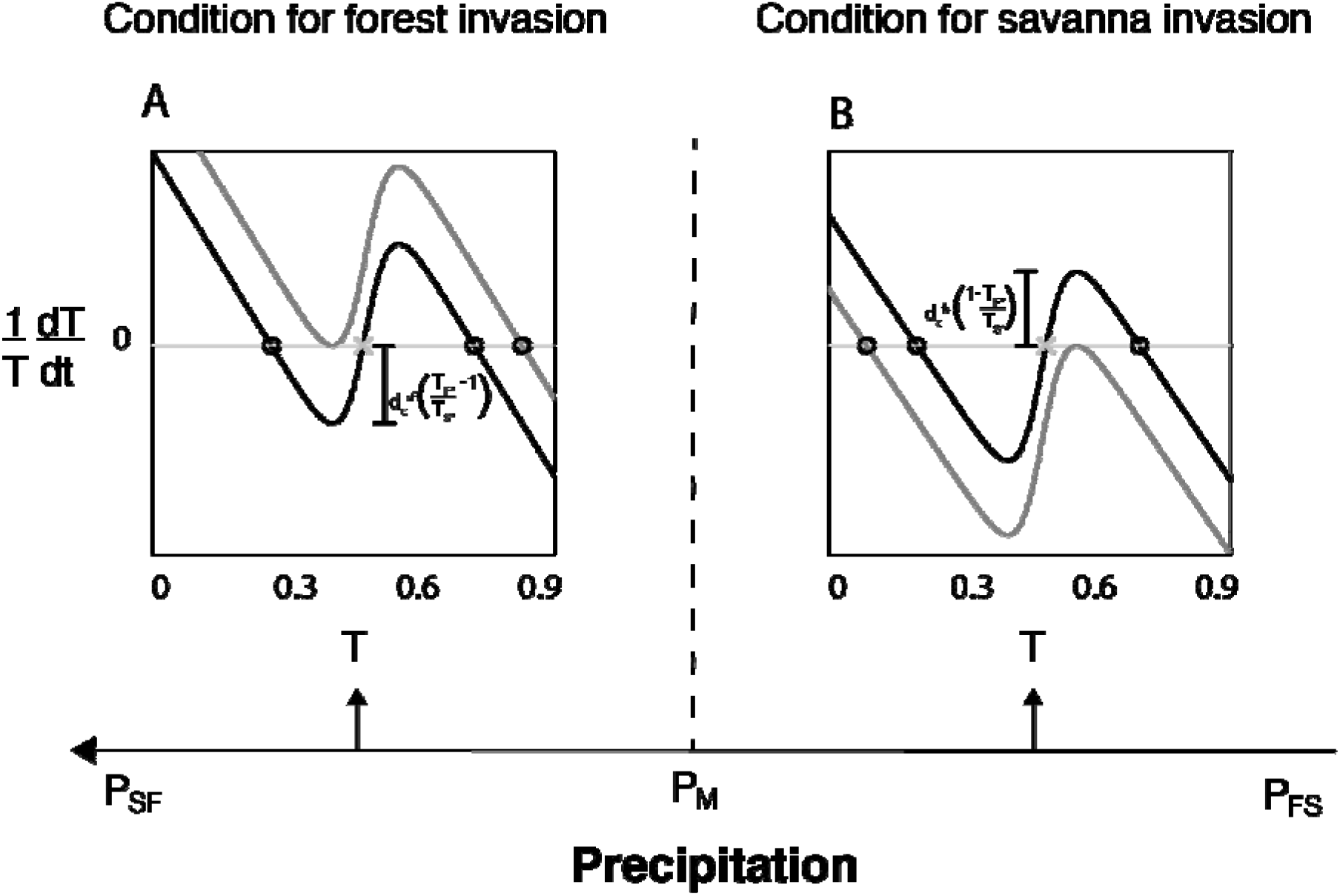
Critical threshold value of dispersal below which the savanna-forest boundary ceases to move despite climatically suitable patches ahead. The black and grey curves in each plot represent the per-capita growth rate (square bracket term in Eqs. 4 and 5) at the barrier when dispersal constantis zero and 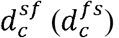, respectively. The length ofI-shaped arrow represents the magnitude of per-capita dispersal flux above which forest fails to invade savanna (A) and vice-versa (B). In particular, when dispersal is below 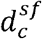, forest fails to invade savanna even though climate favors forest (*i.e.*,*P* > *P*_*M*_)(A). Similarly, when dispersal is below 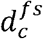, savanna fails to invade forest even though climate favors savanna (*i.e.*,*P* < *P*_*M*_)(B). Thus, in a 1D landscape with a precipitation gradient, the savanna-forest boundarymay stabilize at the dispersal barrier due to fire-vegetation feedback even though it may not have realized its bioclimaticlimit *P*_*M*_. This ecological mechanism—referred to as range pinning—can potentially explain how mesic savannas are maintained in Central Madagascar.

Next, we increase *d*, which graphically corresponds to raising *P*(1 − *T*) − *ϕ* by 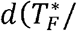 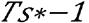. For a low value of *d*, the resulting curve given inside the square brackets again has two stable roots. But if *d* increases past a threshold value, 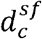, the savanna and unstable roots collide and disappear, and the tree cover jumps to the forest equilibrium (grey curve in Fig. 4A). Thus, for any value of 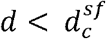, forest fails to invade savanna, *i.e.*, the boundary ceases to move when it encounters the barrier. Intuitively, the forest invasion fails because, in a dispersal limited landscape, fire feedbacks at the boundary savanna patch prevents trees from closing the canopy.

Note that the above graphical argument only provides sufficient conditions on *d* for range pinning. In fact, pinning occurs even when *d* is slightly greater 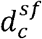 (see Appendix S1). In Fig. S4, we numerically simulate Eq. 4 to obtain necessary and sufficient conditions on *d* for which forest fails to invade savanna, supporting the intuition developed herein. These simulations also reveal that the likelihood of pinning, or the pinning range, increases near *P*_*M*_.

An analogous argument can be constructed to find sufficient conditions on *d* for which savanna fails to invade patch *k* + 1 when *P* < *P*_*M*_. At this patch,

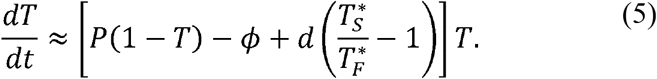

Here again, we lose the subscript *k* + 1 for convenience. As *d* increases, *P*(1 − *T*) − *ϕ* drops by 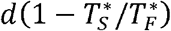. The bracket term has one unstable and two stable roots until a threshold value of *d*, 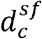, above which the forest and unstable roots collide and disappears, and the tree cover jumps to the savanna equilibrium (Fig. 4B). Thus, in a 1D landscape with precipitation gradient, the boundary can pin at the dispersal barrier, which otherwise would continue until it reaches *P*_*M*_.

The model elaborated above makes three key assumptions that may be violated in Madagascar. First, the model assumes a landscape with one barrier; in reality, landscapes often have multiple barriers. Humphries et al. (2011) showed that multiple barriers have no or little effect on the conditions under which pinning is expected, but may determine where the boundary is pinned. For example, in a landscape with multiple barriers, the boundary will equilibrate at the one that it encounters first, provided all the barriers meet the pinning requirements. Second, we presented pinning conditions at the barrier. It is possible that the boundary may instead pin before reaching the barrier, due to long-range dispersal effects. Using a piecewise linear growth function, we present analytical conditions for pinning for all locations. We find that the boundary may pin at multiple locations near the barrier, but the pinning interval is widest at the dispersal barrier (see Appendix S1 and Fig. S2). Third, a linear barrier in a 2D landscape may have holes (*e.g.*, mountain passes) that may allow the climatically favorable biome state to escape. Using simulations, we show if the width of the pass is small, the boundary may still stabilize at the edges (Fig. S5). Therefore, all scenarios produce results that are qualitatively consistent with our intuition from the 1D stepping-stone model presented above.

#### Large-scale simulations

To check if range pinning is consistent with vegetation patterns in Madagascar, we simulate biome patterns using the 2D calibrated model presented in the previous section. Simulation methods are similar to those described in Goel et al. (2018), but this time, we also include information on the topography. Again, we use a binary dispersal scheme: using the topography map of Madagascar, we code all the patches along the steep slopes (*i.e.*, elevation angle above 3.5 × 10^−3^ rad) of the Central Plateau as dispersal limited (brown patches in Fig. 3B) with *d* equal to 0.1% of *D*. For all the other patches, we used the dispersal rate for the homogeneous stepping-stone model in the previous section (see Fig. S6 in Appendix S1).

The simulation predicts drastically different biome patterns in Madagascar (Fig. 3B) when compared to the 2D diffusion model (Fig. 3A). We find dispersal limitation at the escarpment of the Central Plateau not only maintains mesic savannas in Central Highlands by preventing forest expansion but also maintains rainforests in South-Eastern Madagascar by preventing savanna expansion. Thus, simulation suggests range pinning provides a plausible theoretical explanation for how savannas could have been naturally maintained in mesic regions of Central Highlands.

## Discussion

Here, we develop a stepping-stone model to examine the potential role of dispersal barriers in the maintenance of tropical biomes in Madagascar. Contrary to the predictions of the bioclimatic models (Wuyts et al. 2017), the heterogeneous dispersal model suggests that the savanna-forest boundary in Madagascar is not set by climate (Fig. 3). Instead, topography, in interaction with fire feedbacks, exerts an overriding control in determining the location of the biome boundary. Specifically, the model predicts that the boundary can stabilize at a dispersal barrier despite being far from its bioclimatic limit (Fig. 4). This phenomenon, referred to as range pinning (Keitt et al. 2001), may potentially explain why forests have not expanded into highland savannas, even though these regions are climatically suited to support forest. This mechanism for the maintenance of biomes suggests that savannas in Central Madagascar may not be a product of deforestation.

During the French colonial era, environmentalists in Madagascar perceived fire as a destructive agent that had consumed 90% of the island’s forests (de La Bâthie 1921, Humbert 1927). Post-independence, these views were reinforced by international organizations (Klein 2002), that contributed millions of dollars to fund conservation projects for planting trees and suppressing fires (Scales 2014). In support, the state of Madagascar introduced legislation to discourage burning practices, leading to conflicts between forestry departments and Malagasy farmers who have traditionally relied on fire for pastoral activities (Kull 2000, 2004). Similar misinterpretation of the role of fire in maintaining mesic savannas and domestic conflicts are widespread in tropical regions, including India (Joshi et al. 2018), Brazil (Durigan and Ratter 2016), and West Africa (Fairhead and Leach 1996). These notions have misdirected global conservation efforts in wet savannas that are under threat not only from land-use change (Aleman et al. 2016) but also from ‘reforestation’ projects—such as the Bonn Challenge (WRI 2014)—that are aimed at sequestering carbon to slow global warming (Bastin et al. 2019) and restore ‘lost’ forests that were never forests to begin with. However, a growing theoretical, as well as empirical, literature on tropical plant distributions emphasizes that fire, in combination with other ecological processes, such as dispersal, may play a key role in maintaining overlapping rainfall ranges over which savanna and forest occur.

Broadly, two analytical approaches are used to explain the mismatch between climate and biomes. The mean-field models that ignore dispersal suggest that the climatic mismatch is maintained by fire feedbacks (Beckage et al. 2009, Staver and Levin 2012), which stabilizes initial biome states into local basins of attraction (*e.g.*, in island ecosystems). Meanwhile, 2D diffusion models that incorporate dispersive interactions at the savanna-forest ecotone posit that the climatic mismatch is instead maintained by source-sink dynamics, which is mediated by the geometrical shape of rainfall contours (see Goel et al. 2018, Li et al. 2019, and homogeneous dispersal model above). Most landscapes, however, are dissected by dispersal barriers, such as rivers, plateaus, and mountain ranges, to yield partially disjoint landscape (Rapoport 1982). The stepping stone model, presented here, describes the dynamics of biomes in such a landscape. When the landscape has a high dispersal rate, the pinning region squeezes to *P*_*M*_, and the stepping-stone model behaves like the diffusion model (see Appendix S1). But as dispersal becomes limited, the pinning region widens, and as a result, fire feedbacks may locally stabilize the boundary depending on the patch specific dispersal rate and the climatic conditions. Thus, spatial heterogeneity and local fire feedbacks can maintain savanna in regions where the climate can support forest and vice-versa.

These dynamical properties of the stepping stone model also present a very different view of how biomes will respond to climate change. For starters, range pinning at dispersal barriers may buffer against climate change and introduce spatial lags between bioclimatic and observed spatial limits of biomes (*e.g*., Rupp et al. 2001). Although experimental demonstration of spatial lags is often limited by the time scales of ecological change, paleoecological records corroborate this claim in Madagascar, where the current biome boundary (see Mayaux et al. 2000 and Fig. 1A) is remarkably consistent with Holocene distributions (Adams and Faure 1997b) even though the climate has changed substantially over this time (Buney 1996). There is an important caveat to this prediction, however. If climate change exceeds the pinning threshold (Fig. 4), we should expect to see rapid irreversible biome expansions over large spatial scales. These catastrophic shifts can also incur, for instance, as a result of ill-conceived afforestation plans in Central Highlands (WRI 2014), which alleviates dispersal limitation, allowing rapid forest spread and resulting in loss of both human livelihood and savanna biodiversity.

Conceptually, the discovery of range pinning in Madagascar challenges the hierarchical view of distribution in biogeography (Pearson and Dawson 2003). Bioclimatic models often assume that at the continental scales, the climate is the dominant factor that determines the distribution of biota, whilst the importance of topography (amongst other factors) increases only at more local scales. However, our analysis suggests that synergetic effects of dispersal constraints and non-linear feedbacks can equal or even sometimes outweigh the contribution of the climate in explaining tropical plant distributions. More broadly, since negative growth rate at low abundance is a ubiquitous in plant and animal populations (Allee 1927, Allee et al. 1949), even partially porous barriers can, in effect, become absolute barriers and stabilize geographical ranges over wide climatic conditions.

## Supporting information

Metadata S1

## Acknowledgments

We thank Staver Lab members, particularly Madelon Case and Joshua Daskin, for their comments. We also thank Arun Chavan, Stephen Stearns, Timothy Keitt, Simon Stump, and Fiona MacNeill for their feedback on writing. NG and ACS were supported by NSF DMS-1615531 to ACS and EVV by NSF DMS-1419047. NG acknowledges support from the Yale Institute for Biospheric Studies. NG and ACS designed the study. NG and JCA gathered the paleoecological data. NG and EVV designed the model. NG and ACS wrote the paper with feedback from other authors. Authors declare no competing interests.

## References

Adams, J. M., and H. Faure. 1997a. Preliminary vegetation maps of the world since the last glacial maximum: An aid to archaeological understanding. Journal of Archaeological Science 24:623–647.

Adams, J. M., and H. Faure. 1997b. QEN members. Review and atlas of palaeovegetation: preliminary land ecosystem maps of the world since the Last Glacial Maximum. Oak Ridge National Laboratory, TN. Page https://www.esd.ornl.gov/projects/gen/nerc.html.

Aleman, J. C., O. Blarquez, and C. A. Staver. 2016. Land-use change outweighs projected effects of changing rainfall on tree cover in sub-Saharan Africa. Global Change Biology 22:3013–3025.

Allee, W. C. 1927. Animal aggregations. The Quarterly Review of Biology 2:367–398.

Allee, W. C., O. Park, A. E. Emerson, T. Park, and K. P. Schmidt. 1949. Principles of animal ecology. Saunders Company Philadelphia, Pennsylvania, USA.

Archibald, S., D. P. Roy, V. Wilgen, W. Brian, and R. J. Scholes. 2009. What limits fire? An examination of drivers of burnt area in Southern Africa. Global Change Biology 15:613–630.

Bastin, J.-F., Y. Finegold, C. Garcia, D. Mollicone, M. Rezende, D. Routh, C. M. Zohner, and T. W. Crowther. 2019. The global tree restoration potential. Science 365:76–79.

Beckage, B., W. J. Platt, and L. J. Gross. 2009. Vegetation, fire, and feedbacks: a disturbance-mediated model of savannas. The American Naturalist 174:805–818.

Bond, W. J., G. F. Midgley, F. I. Woodward, M. T. Hoffman, and R. M. Cowling. 2003. What controls South African vegetation—climate or fire? South African Journal of Botany 69:79–91.

Bond, W. J., J. A. Silander Jr, J. Ranaivonasy, and J. Ratsirarson. 2008. The antiquity of Madagascar’s grasslands and the rise of C4 grassy biomes. Journal of Biogeography 35:1743–1758.

Bond, W. J., F. I. Woodward, and G. F. Midgley. 2005. The global distribution of ecosystems in a world without fire. New phytologist 165:525–537.

Bowman, D. M., and G. L. Perry. 2017. Soil or fire: what causes treeless sedgelands in Tasmanian wet forests? Plant and soil 420:1–18.

Buney, D. 1996. Climate change and fire ecology as factors in the Quaternary biogeography. Biogéographie de Madagascar:49–58.

de La Bâthie, H. P. 1921. La végétation malgache. Musée Colonial.

Durigan, G., and J. A. Ratter. 2016. The need for a consistent fire policy for Cerrado conservation. Journal of Applied Ecology 53:11–15.

Fairhead, J., and M. Leach. 1996. Misreading the African landscape: society and ecology in a forest-savanna mosaic. Cambridge University Press.

Gasse, F., and E. Van Campo. 1998. A 40,000-yr pollen and diatom record from Lake Tritrivakely, Madagascar, in the southern tropics. Quaternary Research 49:299–311.

Gaston, K. J. 2003. The structure and dynamics of geographic ranges. Oxford University Press on Demand.

Goel, N., V. Guttal, S. Levin, and C. Staver. 2018. Dispersal increases the resilience of tropical savanna and forest distributions. bioRxiv:476184.

Goldberg, E. E., and R. Lande. 2007. Species’ borders and dispersal barriers. The American Naturalist 170:297–304.

Green, G. M., and R. W. Sussman. 1990. Deforestation history of the eastern rain forests of madagascar from satellite images. Science 248:212–215.

Hansford, J., P. C. Wright, A. Rasoamiaramanana, V. R. Perez, L. R. Godfrey, D. Errickson, T. Thompson, and S. T. Turvey. 2018. Early Holocene human presence in Madagascar evidenced by exploitation of avian megafauna. Science Advances 4:eaat6925.

Hennenberg, K. J., F. Fischer, K. Kouadio, D. Goetze, B. Orthmann, K. E. Linsenmair, F. Jeltsch, and S. Porembski. 2006. Phytomass and fire occurrence along forest--savanna transects in the Como’e National Park, Ivory Coast. Journal of Tropical Ecology 22:303–311.

Higgins, S. I., W. J. Bond, and W. S. W. Trollope. 2000. Fire, resprouting and variability: a recipe for grass-tree coexistence in savanna. Journal of Ecology 88:213–229.

Hirota, M., M. Holmgren, E. H. Van Nes, and M. Scheffer. 2011. Global resilience of tropical forest and savanna to critical transitions. Science 334:232–235.

Hoffman, A., H. Hupkes, and E. Van Vleck. 2017. Entire solutions for bistable lattice differential equations with obstacles. American Mathematical Society.

Holdridge, L. R. 1947. Determination of World Plant Formations From Simple Climatic Data. Science 105:367–368.

Huffman, G. J., and D. T. Bolvin. 2013. TRMM and other data precipitation data set documentation. NASA, Greenbelt, USA 28:1.

Humbert, H. 1927. Destruction d’unefloreinsulaire par le feu.

Humbert, H., and G. Cours Darne. 1965. Carte Internationale du Tapis Végétal: Madagascar, 1∶ 1,000,000. French Institute of Pondichery, Toulouse.

Humphries, A. R., B. E. Moore, and E. S. Van Vleck. 2011. Front Solutions for Bistable Differential-Difference Equations with Inhomogeneous Diffusion. SIAM Journal on Applied Mathematics 71:1374–1400.

Jarvis, A., H. I. Reuter, A. Nelson, and E. Guevara. 2008. Hole-filled SRTM for the globe Version 4. available from the CGIAR-CSI SRTM 90m Database (http://srtm.csi.cgiar.org).

Joshi, A. A., M. Sankaran, and J. Ratnam. 2018. ‘Foresting’ the grassland: Historical management legacies in forest-grassland mosaics in southern India, and lessons for the conservation of tropical grassy biomes. Biological conservation 224:144–152.

Keitt, T. H., M. A. Lewis, and R. D. Holt. 2001. Allee effects, invasion pinning, and species’ borders. The American Naturalist 157:203–216.

Klein, J. 2002. Deforestation in the Madagascar highlands--establishedtruth’and scientific uncertainty. GeoJournal 56:191–199.

Koechlin, J. 1972. Flora and vegetation of Madagascar. Pages 145–190 Biogeography and ecology in Madagascar. Springer.

Kull, C. A. 2000. Deforestation, erosion, and fire: degradation myths in the environmental history of Madagascar. Environment and History 6:423–450.

Kull, C. A. 2004. Isle of fire: the political ecology of landscape burning in Madagascar. University of Chicago Press.

Lehmann, C. E., S. A. Archibald, W. A. Hoffmann, and W. J. Bond. 2011. Deciphering the distribution of the savanna biome. New phytologist 191:197–209.

Levin, S. A. 1992. The problem of pattern and scale in ecology: the Robert H. MacArthur award lecture. Ecology 73:1943–1967.

Li, Q., A. C. Staver, E. Weinan, and S. A. Levin. 2019. Spatial feedbacks and the dynamics of savanna and forest. Theoretical Ecology:1–26.

Lloyd, J., M. I. Bird, L. Vellen, A. C. Miranda, E. M. Veenendaal, G. Djagbletey, H. S. Miranda, G. Cook, and G. D. Farquhar. 2008. Contributions of woody and herbaceous vegetation to tropical savanna ecosystem productivity: a quasi-global estimate. Tree physiology 28:451–468.

Marsden, S. S. A. J. E., L. S. S. Wiggins, L. Glass, R. V. Kohn, and S. S. Sastry. 1993. Interdisciplinary Applied Mathematics. Springer.

Mayaux, P., V. Gond, and E. Bartholome. 2000. A near-real time forest-cover map of Madagascar derived from SPOT-4 VEGETATION data. International Journal of Remote Sensing 21:3139–3144.

McKean, H. P. 1970. Nagumo’s equation. Advances in mathematics 4:209–223.

Menaut, J. 1983. The vegetation of African savannas. Ecosystems of the World.

Moreira, A. G. 2000. Effects of fire protection on savanna structure in Central Brazil. Journal of Biogeography 27:1021–1029.

Murray, J. D. 2001. Mathematical Biology. II Spatial Models and Biomedical Applications. Springer-Verlag New York Incorporated.

Okubo, A. 1986. Dynamical aspects of animal grouping: swarms, schools, flocks, and herds. Advances in biophysics 22:1–94.

Pearson, R. G., and T. P. Dawson. 2003. Predicting the impacts of climate change on the distribution of species: are bioclimate envelope models useful? Global Ecology and Biogeography 12:361–371.

Prior, L. D., R. J. Williams, and D. M. J. S. Bowman. 2010. Experimental evidence that fire causes a tree recruitment bottleneck in an Australian tropical savanna. Journal of Tropical Ecology 26:595–603.

Pueyo, S., P. M. de Alencastro Graca, R. I. Barbosa, R. Cots, E. Cardona, and P. M. Fearnside. 2010. Testing for criticality in ecosystem dynamics: the case of Amazonian rainforest and savanna fire. Ecology Letters 13:793–802.

Purata, S. E. 1986. Floristic and structural changes during old-field succession in the Mexican tropics in relation to site history and species availability. Journal of Tropical Ecology 2:257–276.

Rapoport, E. H. 1982. Areography: geographical strategies of species. Elsevier.

Rupp, T. S., F. S. Chapin, and A. M. Starfield. 2001. Modeling the influence of topographic barriers on treeline advance at the forest-tundra ecotone in northwestern Alaska. Climatic change 48:399–416.

Scales, I. R. 2014. Conservation and environmental management in madagascar. Routledge.

Skellam, J. G. 1951. Random dispersal in theoretical populations. Biometrika 38:196–218.

Staal, A., S. C. Dekker, C. Xu, and E. H. van Nes. 2016. Bistability, Spatial Interaction, and the Distribution of Tropical Forests and Savannas. Ecosystems 19:1080–1091.

Staver, A. C., S. Archibald, and S. Levin. 2011a. Tree cover in sub-Saharan Africa: Rainfall and fire constrain forest and savanna as alternative stable states. Ecology 92:1063–1072.

Staver, A. C., S. Archibald, and S. A. Levin. 2011b. The Global Extent and Determinants of Savanna and Forest as Alternative Biome States. Science 334:230–232.

Staver, A. C., and S. A. Levin. 2012. Integrating theoretical climate and fire effects on savanna and forest systems. The American Naturalist 180:211–224.

Swaine, M. D., W. D. Hawthorne, and T. K. Orgle. 1992. The Effects of Fire Exclusion on Savanna Vegetation at Kpong, Ghana. Biotropica 24:166–172.

Taylor, C. M., and A. Hastings. 2005. Allee effects in biological invasions. Ecology Letters 8:895–908.

Touboul, J. D., A. C. Staver, and S. A. Levin. 2018. On the complex dynamics of savanna landscapes. Proceedings of the National Academy of Sciences 115:E1336–E1345.

Trapnell, C. G. 1959. Ecological results of woodland and burning experiments in Northern Rhodisia. The Journal of Ecology:129–168.

Veenendaal, E. M., M. Torello-Raventos, H. S. Miranda, N. M. Sato, I. Oliveras, F. van Langevelde, G. P. Asner, and J. Lloyd. 2018. On the relationship between fire regime and vegetation structure in the tropics. New phytologist 218:153–166.

Vorontsova, M. S., G. Besnard, F. Forest, P. Malakasi, J. Moat, W. D. Clayton, P. Ficinski, G. M. Savva, O. P. Nanjarisoa, and J. Razanatsoa. 2016. Madagascar’s grasses and grasslands: anthropogenic or natural? Proceedings of the Royal Society B: Biological Sciences 283:20152262.

Wang, C. H., S. Matin, A. B. George, and K. S. Korolev. 2019. Pinned, locked, pushed, and pulled traveling waves in structured environments. Theoretical population biology 127:102–119.

Woodward, F. I. 1987. Climate and plant distribution. Cambridge University Press.

WRI. 2014. Atlas of Forest and Landscape Restoration Opportunities. World Resources Institute: Washington, DC. Atlas of Forest and Landscape Restoration Opportunities. World Resource Institute.

Wuyts, B., A. R. Champneys, and J. I. House. 2017. Amazonian forest-savanna bistability and human impact. Nature Communications 8:15519.

Wuyts, B., A. R. Champneys, N. Verschueren, and J. I. House. 2019. Tropical tree cover in a heterogeneous environment: A reaction-diffusion model. PloS one 14:e0218151.

Yoder, A. D., C. R. Campbell, M. B. Blanco, M. Dos Reis, J. U. Ganzhorn, S. M. Goodman, K. E. Hunnicutt, P. A. Larsen, P. M. Kappeler, R. M. Rasoloarison, J. M. Ralison, D. L. Swofford, and D. W. Weisrock. 2016. Geogenetic patterns in mouse lemurs (genus Microcebus) reveal the ghosts of Madagascar’s forests past. Proceedings of the National Academy of Sciences 113:8049–8056.

