## Appendix S1 for "Dispersal limitation and fire feedbacks maintain mesic savannas in Madagascar"

Here, we provide details of the empirical and mathematical analysis underlying the main text. We organize our supplementary information as follows. First, we provide details on how we collected and reanalyzed paleoecological data from published sources. In the second section, we present analytical solutions for range pinning for a piecewise linear growth function. Lastly, we describe numerical methods to simulate biome distribution in Madagascar with and without dispersal barriers.

### Paleo analysis

We reconstruct the historical vegetation patterns in Madagascar using three non-climatic proxies: (*i*) plant pollen, (*ii*) $\delta^{13}$C, and (*iii*) charcoal. Plant pollen is used to identify the main plant taxa and, therefore, the biome type in the source area of the sedimentary archive. Sites with a higher percentage of Gramineae represented in the pollen assemblage are classified as savanna [for more details see Gasse et al. (1994)]. The $\delta^{13}$C value of an archive can be used to estimate the relative abundance of plants that photosynthesize using $C_{4}$ and $C_{3}$ pathways. Since grasses and trees in tropics use $C_{4}$ and $C_{3}$ pathway for photosynthesis, respectively, the $\delta^{13}$C value can be used to distinguish savanna and forests (Boutton et al. 1999, Aleman et al. 2012). Finally, we use charcoal particles recorded in the sediments as a proxy for fire activity (Power et al. 2008), characteristic of savanna vegetation. Please note that we reconstruct historical biome patterns using paleo proxies that depend on regional functional traits of plants (like flammability and photosynthetic pathway) that although may be correlated with climate, are not strictly determined by it.

In total, we gathered 19 published records (see Metadata S1): 13 with combined pollen and charcoal proxies, 1 with only pollen, 4 with only charcoal, and 1 with $\delta^{13}$C analysis of speleothems. Of the 19 sites, we excluded the seven sites because of poor age-depth models and incomplete pollen analysis (#12-17), with the remaining 11 used in Fig. 1A. Table S1 shows the time for the increase in the percentage of Gramineae pollen and charcoal flux (characteristic of savanna vegetation) at various locations in Madagascar. When dates in the original papers were not calibrated, we used SHCal13 calibration curve from Hogg et al. (2013) to convert radiocarbon dates to calendar years BP (before present, the present being 1950 by convention). Here, we present conservative estimates for the presence of savanna vegetation (column 6). In fact, many sites show fire activity much before the date we classify them as savanna sites (see column 7 in Table S1).

### Piecewise linear growth function

In the main text, we present a graphical approach to understanding savanna-forest boundary dynamics in homogeneous and heterogenous dispersal environment. However, that graphical approach in the main text is limited when compared to an analytical approach because the latter captures the dynamics of the system for all parameter values and not just for a select few. This allows us to gain deeper and general insights that are not otherwise possible. Unfortunately, obtaining an analytical solution for Eq. 2 is hard because of the non-linear terms in Eq. 1.

Here, we present a workaround that considers a growth function that is both bistable and linear:

$$f\left( T,P \right)=-T+ \Theta(T-P),$$

(S1)

where $\Theta(x)$ is a step function, *i.e.*, $\Theta(x\leq0)=0$ and $\Theta(x>0)=1$ (Fig. S1). Note, when $T\leq P$, $T=0$ is stable, and when $T>P$, $T=1$ is stable. We assume that *zero* (*one*) root represents the savanna (forest) state. Here again, we assume $T+G=1$. Thus, when the system is in the savanna (forest) attractor, grass density is *unity* (*zero*). In the mathematical literature, Eq. S1 is referred to as McKean’s caricature of the cubic. Although McKean’s caricature is not biologically motivated, it captures the bistability of tropical biomes that is central to explaining the results of the paper.

Below we present the analytical results for both 1D homogenous and 1D heterogeneous dispersal model. Please refer to Fáth (1998) and Humphries et al. (2011) for mathematical details.

#### Homogeneous dispersal model

Recall, in a homogeneous dispersal model, we assume $D_{i} = D\gg1$ for all $i$. For this dispersal scheme, we can approximate Eq. 2 as a reaction-diffusion model (Eq. 3). This diffusion model can be analytically solved using Fourier Transform (Fáth 1998) to obtain the speed ($v$) of the savanna-forest boundary as a function of precipitation $P$ and dispersal constant $D$:

$$v={(P}_{M}-P)\sqrt{\frac{4D}{P\left( 1-P \right)}},$$

(S2)

where $P_{M}=0.5$ is the Maxwell precipitation. Note, when $P>P_{M}$, forest invades savanna ($v<0$), and, conversely, when $P<P_{M}$ savanna invades forest ($v>0$) (left plot in Fig. S2). The boundary is stationary ($v=0$) at a unique precipitation value, $P_{M}$. Thus, in a landscape with a linear gradient of precipitation, the boundary equilibrates at the spatial location that receives $P=P_{M}$, where $P_{M}$ can be interpreted as the bioclimatic limit of savanna and forest biomes.

#### Heterogeneous dispersal model

Recall, in a heterogeneous dispersal model, we assume a binary dispersal scheme: $D_{k} = d$ and $D_{i\neq k} = D$ such that $D\gg d$, where $D_{k}$ is the dispersal rate between patches $k$ and $k+1$. Without loss of generality, we assume $k=0$. Using Jacobi Operator Theory, this model can be analytically solved to obtain the equilibrium position of the boundary as a function of background dispersal rate $D$, dispersal rate at the barrier $d$, precipitation $P$, and distance from the barrier $l=i-k$ (Humphries et al. 2011).

We find that the boundary stabilizes between patches $0$ and $1$ if precipitation lies in the following interval,

$$\left[ \frac{\tau}{\lambda+2\tau-1},\frac{\lambda+\tau-1}{\lambda+2\tau-1} \right]=[a,b],$$

(S3)

where $\lambda=1+\frac{1}{2D}(1+\sqrt{1+4D})$ and $\tau=d/D$ (Fig. S3). This precipitation interval is called the interval of propagation failure. Note, for fixed $d$, if $P_{M}<P<b$, forest fails to invade savanna, and conversely, if $P_{M}>P>a$, savanna fails to invade forest. In both cases, the invasion fails despite climatically suitable patches ahead. For consistency with the main text, we invert this interval to find conditions on $d$ for which invasion fails (Fig. S3). Here too, for fixed $P>P_{M}$, if $d<d_{sf}$, forest fails to invade savanna. Conversely, for fixed $P<P_{M}$, if $d<d_{fs}$, savanna fails to invade forest.

The boundary can also stabilize at $i<k$, if $P$ lies within

|  | $\left[ \frac{1}{\lambda+1}\left\{ 1-\lambda^{2l}\left( a-\lambda b \right) \right\}, \frac{\lambda}{\lambda+1}\left\{ 1-\lambda^{2l}\left( a-\lambda b \right) \right\} \right],$ | (S4) |
| --- | --- | --- |

and at $i>k$, if $P$ lies within

|  | $\left[ \frac{1}{\lambda+1}\left\{ 1-\lambda^{1-2l}\left( a-\lambda b \right) \right\}, \frac{\lambda}{\lambda+1}\left\{ 1-\lambda^{-\left( 1+2l \right)}\left( a-\lambda b \right) \right\} \right].$ | (S5) |
| --- | --- | --- |

These calculations reproduce several results that we presented using a graphical approach in the main text. But these calculations also offer additions insights. Below we discuss some of these analytical insights.

First, if there is no dispersal barrier, *i.e.*, $d=D\gg1$, the interval of propagation failure approaches [$P_{M}^{-} ,P_{M}^{+}$] for all patches [substitute $\tau=\lambda=1$ in equation S3-S5]. In other words, the heterogeneous dispersal model behaves like a reaction-diffusion described above. Second, if $d$ is low, the boundary may stabilize at the barrier if the precipitation meets the condition in Eq. S3. Graphically, this corresponds to the situation where $d$ and $P$ fall in the grey region in Fig. S3. Moreover, the interval of propagation failure at the barrier widens as $d$ decreases. When $d=0$, the interval is equal to the bistable region [$0,1$], i.e., the dynamics at the boundary savanna and forest patches become decoupled and is governed by independent mean-field growth function in Eq. S1. Third, for fixed $d$, the interval of propagation failure is widest at the barrier and decreases as one moves far away from the barrier (right plot in Fig. S2). Finally, when the boundary is sufficiently far away from the barrier, the interval of propagation failure converges to [$P_{M}^{-} ,P_{M}^{+}$] (right plot in Fig. S2). In other words, the heterogeneous dispersal model behaves like a homogeneous reaction-diffusion when the boundary is far away from the barrier.

### Large-scale simulations

Here, we describe methods for simulating the distribution of tropical biomes in Madagascar for the two dispersal schemes considered in the main text using R version 3.5.5.

#### Datasets

For our simulations, we used three raster datasets: (*i*) biome distributions, (*ii*) rainfall patterns, and (*iii*) a topography map of Madagascar. All raster datasets were transformed to have the same extent [${43}^{o}$E-${51}^{o}$E and ${26}^{o}$S-${12}^{o}$S] and resolution [${0.0625}^{o} x {0.0625}^{o}$(~ $6.5$ km $x$ $6.5$ km)]. Furthermore, all the pixels corresponding to water bodies were excluded from the analysis.

**Vegetation*.***

The forest-cover map used in this paper was derived using SPOT-4 VEGETATION data from October 1998 to September 1999 (Mayaux et al. 2000). We used this dataset because the authors provided an objective method to identify deforested areas in Madagascar. The proposed forest extent (present-day forest extent + deforested region) in their paper is consistent with the aerial photos of vegetation cover from 1950 (Green and Sussman 1990), published extent of forest (Koechlin 1972, Bond et al. 2008) on the island, and the paleo analysis (Fig. 1A and Table S1).

**Rainfall.**

We used the mean annual precipitation (MAP) in Madagascar from 2000 to 2010 derived from Tropical Rainfall Measuring Mission (Huffman and Bolvin 2013).

**Topography.**

We used Shuttle Radar Topographic Mission 90 m Digital Elevation Data [SRTM 90m DEM] produced by NASA (Jarvis et al. 2008). We calculated the elevation gradient from the elevation raster using $\sqrt{\partial^{2}/\partial x^{2}+\partial^{2}/\partial y^{2}}$operator. Intuitively, the gradient operator corresponds to the slope of the landscape which for small slope is equal to the angle itself, *i.e.*, $\tan\theta= \theta$ if $\theta\ll1$.

#### Large-scale simulations

Although in the main text, we have presented analytical results for 1D models, for large-scale simulations, we use the 2D model for both dispersal schemes. Here, we first describe the numerical steps that are common for both dispersal schemes, followed by steps that are unique to each scheme. Our method closely follows Goel et al. (2018).

For both dispersal schemes, we use the imperfect bifurcation model,

$$f\left( T,P \right)= -P+3T-T^{3},$$

(S6)

to simulate biome distributions. Using basic algebra, we can show that Eq. S6 has three equilibrium points for$P_{FS}<P<P_{SF}$ (bistable region). The positive (negative) stable root represents forest (savanna). We can also show that $P_{FS}=-2$, $P_{SF}=2$, and $P_{M}=0$.

To simulate vegetation distribution in Madagascar, we first rescale the precipitation raster $P_{old}$ using the following relation: $P_{new}= (P_{old}-P_{M})/350$. We used this re-scaling relationship because we know from previous studies that $P_{FS}$, $P_{SF}$, and $P_{M}$ are approximately 800, 2200, and 1500 mm MAP, respectively. By substituting these values in the rescaling relationship, we recover the critical precipitation and Maxwell precipitation values of the imperfect bifurcation model. Next, we initialize both simulations with 1900 forest cover (present-day forest extent + deforested region) from (Mayaux et al. 2000). To do this, we initialize all patch location corresponding to the forest patches as +2 (positive root) and savanna patches as -2 (negative root).

**Homogeneous dispersal model.**

For the homogeneous dispersal model, we use 2D version of Eq. 2 with $D_{i}= 34$ for all $i$, and $P_{M} = 1538$ mm MAP. These parameter values were obtained from Goel et al. (2018). To implement this dispersal scheme, we use the following algorithm:

$$T_{i,j}^{t+1}=T_{i,j}^{t}+\Delta t\left[ f\left( T_{i,j}^{t},P_{new} \right)+D\left( T_{i-1,j}^{t}+T_{i+1,j}^{t}+T_{i,j-1}^{t}+T_{i,j+1}^{t}-4T_{i,j}^{t} \right) \right],$$

(S7)

where $i$ and $j$ represent the cell coordinates. We truncated the simulations when $\sum_{i,j} |T_{i,j}^{t+1}-T_{i,j}^{t}|<0.1$.

**Heterogeneous dispersal model.**

To study the effects of topography, we simulated the distribution of savanna and forest following the same approach as in the homogeneous dispersal model, except here we implement a heterogeneous dispersal scheme. We declared all the pixels with an elevation gradient above a threshold value $E_{th}=3.5$ x ${10}^{-3}$ rads with low dispersal coefficient, $d= {10}^{-1}$% of $D$. For all other pixels, we used the same value of $D$ as in homogeneous dispersal model. To implement this dispersal scheme, we use the following algorithm:

$$T_{i,j}^{t+1}=T_{i,j}^{t}+\Delta t\left[ f\left( T_{i,j}^{t},P_{new} \right)+D_{i,j}\left[ T_{i+1,j}^{t}-{2T}_{i,j}^{t}+T_{i,j+1}^{t} \right]+D_{i-1,j}\left[ T_{i-1,j}^{t}-T_{i,j}^{t} \right]+D_{i,j-1}[T_{i,j-1}^{t}-T_{j}^{t}] \right],$$

(S8)

where $i$ and $j$ represent the cell coordinates. Here again, we truncated the simulations when $\sum_{i,j} |T_{i,j}^{t+1}-T_{i,j}^{t}|<0.1$.

Since $E_{th}$ and $d$ were chosen arbitrarily in the above analysis, we simulated biome patterns for a wide range of parameter values of $E_{th}$ (from $2.4$ x ${10}^{-3}$ to $4$ x ${10}^{-3}$ rads) and $d$ (from ${10}^{-2}$% to $10$% of $D$) (Fig. S6). Sensitivity analysis shows that the savanna-forest boundary gets pinned at the eastern edge of the Central Plateau for a wide range of parameter values of $E_{th}$ and $d$, suggesting results are robust to variations in the parameter values.

### Tables

**Table S1:** Reconstructed vegetation patterns in Madagascar over last 17 kyrs using non-climatic proxies—pollen, charcoal, and $\delta^{13}$C (speleothems)—from published literature (see Fig. 1A in the main text). Although the time period of most proxies overlaps with human presence in Madagascar, four of them [marked with *; #2, 3, 4, and 11] predate human use of fire in Madagascar (Burney 1999). Of these, the savanna record from Lake Tritrivakely [#4] in Central Madagascar predates human arrival by 6.4 kyrs.

| # | Sites | Lat | Lon | Proxy | Date for savanna biome | Date since charcoal accumulation | References |
| --- | --- | --- | --- | --- | --- | --- | --- |
| 1 | Anjohibe caves | -15.54 | 46.89 | $\delta^{13}$C | 1,060 BP | - | (Burns et al. 2016) |
| 2* | Ambolistra/ Lake Andolonomby | -23.05 | 43.6 | Pollen & Charcoal | 3,050 BP | 5,600 BP | (Burney 1993, Burney et al. 2003) |
| 3* | Lake Mitsinjo | -16.03 | 45.85 | Pollen & Charcoal | 3,760 BP | 3,760 BP | (Matsumotot and Burney 1994, Wright et al. 1996) |
| 4* | Lake Tritrivakely | -19.78 | 46.92 | Pollen & Charcoal | 17,000 BP | 12,000 BP | (Gasse et al. 1994) |
| 5 | Lake Kavitaha | -19.04 | 46.74 | Pollen & Charcoal | 1,050 BP | 1,370 BP | (Burney 1987) |
| 6 | Lake Komango | -19.16 | 44.81 | Charcoal | 1,580 BP | 3,100 BP | (Burney 1999) |
| 7 | Benavony | -13.71 | 48.49 | Charcoal | 650 BP | 4,400 BP | (Burney 1999) |
| 8 | Belo-sur-mer | -20.73 | 44.02 | Charcoal | 1680 BP | 2,030 BP | (Burney 1999) |
| 9 | Amparihibe | -13.3 | 48.22 | Charcoal | 990 BP | 1,920 BP | (Burney 1999) |
| 10 | Mandena (matrix) | -24.93 | 47.01 | Pollen & Charcoal | 1,400 & 850 BP | 5,600 BP | (Virah-Sawmy et al. 2009) |
| 11* | Ste-Luce (matrix) | -24.78 | 47.16 | Pollen & Charcoal | 5,800 BP | 5,800 BP | (Virah-Sawmy et al. 2009) |

**Table S2:** Symbology

| Symbol | Ecological interpretation |
| --- | --- |
| $T_{i}$ | Forest tree density in patch $i$ [state variable] |
| $G_{i}=1-T_{i}$ | Grass density in patch $i$ [state variable] |
| $P$ | Precipitation [control variable] |
| $T_{F}^{*}(T_{S}^{*})$ | Equilibrium tree cover corresponding to the stable forest (savanna) state |
| $T_{critical}^{*}$ | Unstable equilibrium that separates stable forest ($T_{F}^{*}$) and savanna ($T_{S}^{*}$) states |
| $P_{FS} (P_{SF})$ | Critical precipitation value corresponding to forest to savanna transition (and vice-versa) |
| $\phi$ | Per-capita forest tree mortality rate |
| $D_{i}(D_{i,j})$ | Dispersal constant corresponding to patch $i$ in 1D landscape [$(i,j$) in 2D landscape] |
| $\Phi_{i}(\Phi_{i,j})$ | Dispersal flux at patch $i$ 1D landscape [$(i,j$) in 2D landscape] |
| $\Phi_{x}(\Phi_{x,y})$ | Dispersal flux at location $x$ in a 1D landscape [($x,y$) in a 2D landscape] |
| $P_{M}$ | Maxwell precipitation |
| $P_{Mc}$ | Maxwell precipitation contour |
| $d$ | Dispersal constant at the dispersal barrier |
| $D$ | Dispersal constant everywhere except at the barrier |
| $d_{c}^{sf}(d_{c}^{fs})$ | Thereshold value of $d$ below which forest (savanna) fails to invade savanna (forest) |

### Figures

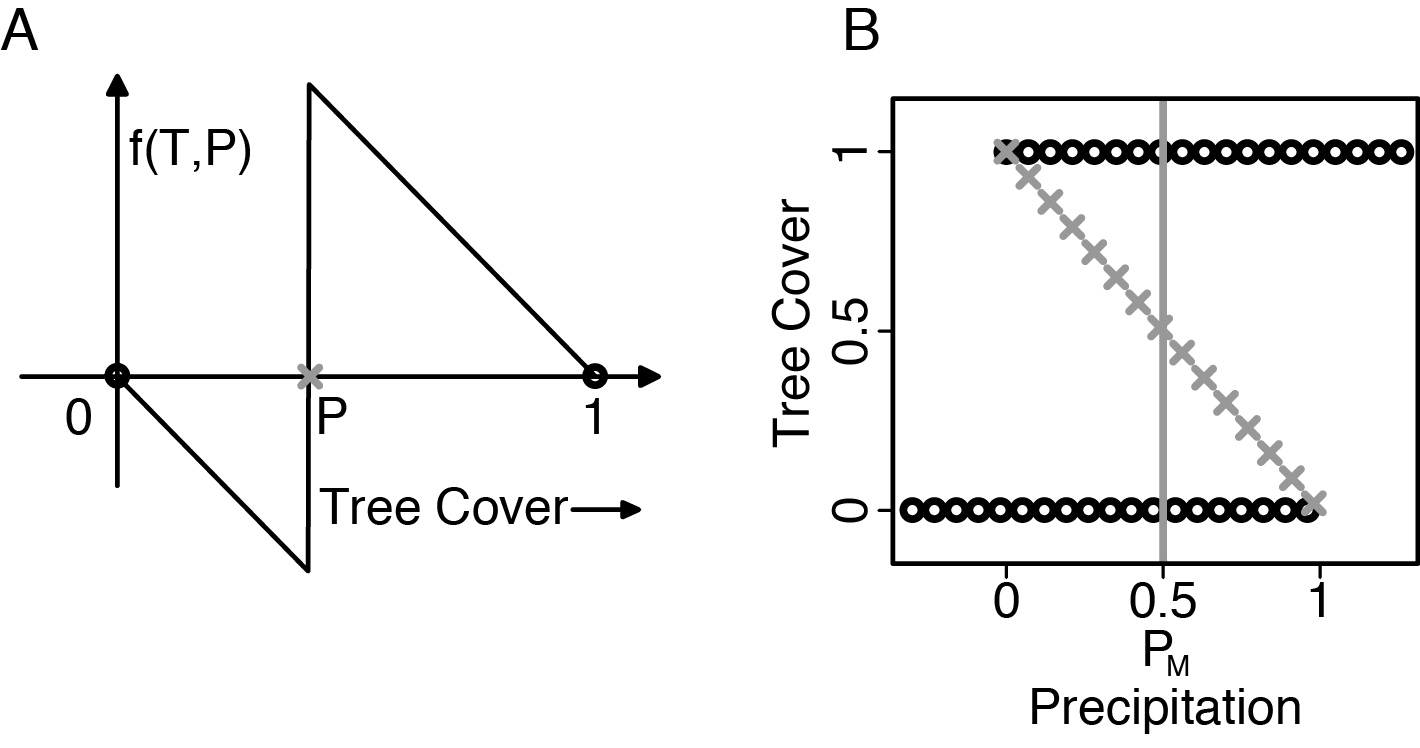

**Figure S1:** Growth rate (A) and bifurcation diagram (B) of piecewise linear growth function (Eq. S1; McKean’s caricature of the cubic). This growth function is both linear (A) and bistable (B). When $T<P$, savanna is stable, and conversely, when $T>P$ forest is stable. The vertical grey line is the $P_{M}=0.5$. The black circles and grey crosses indicate stable equilibrium points and separatrix of the system.

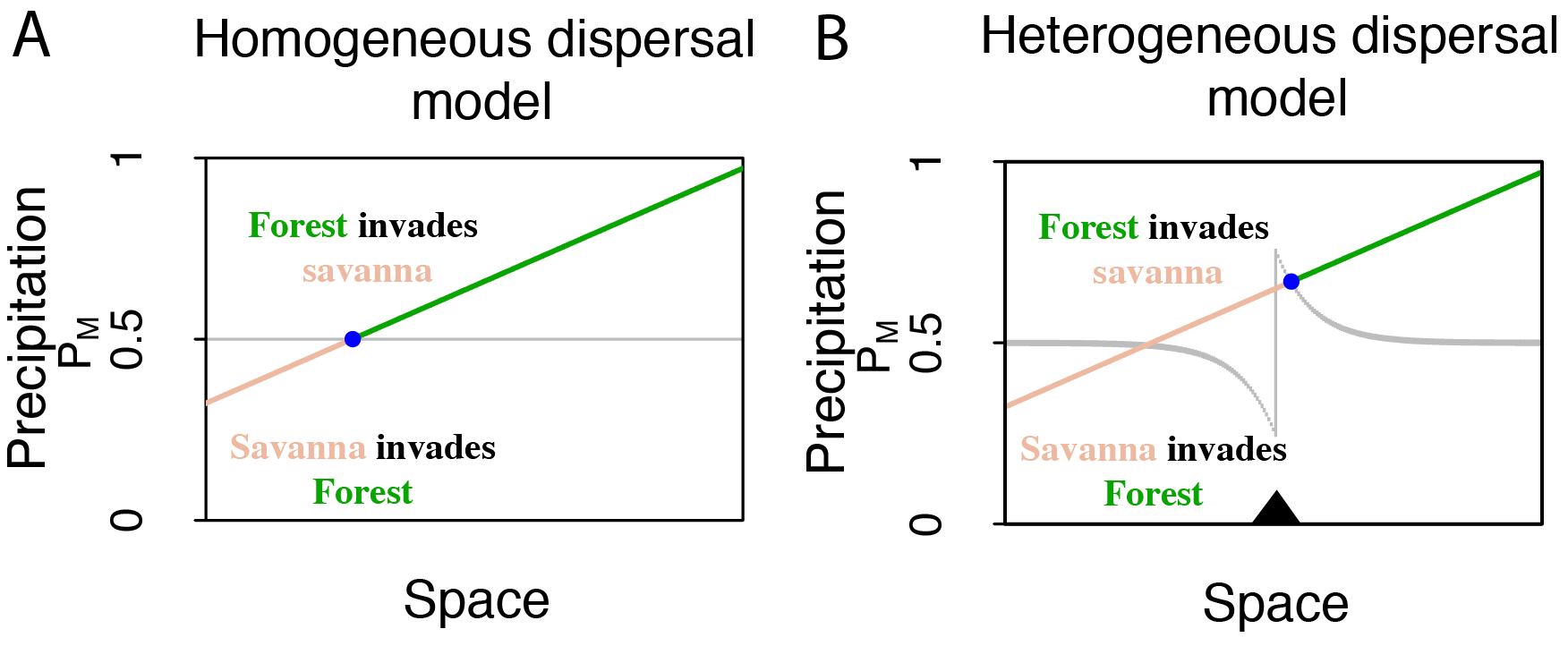

**Figure S2:** Phase plot of the direction of movement of the savanna-forest boundary as a function of precipitation and spatial coordinate in 1D for Eq. 1 with piece wise linear growth function in Eq. S1. The grey line in the plots (A) and (B) represent the precipitation value(s) for which $v=0$. Above the grey region, forest invades savanna, and vice-versa. The triangle in the plot (B) represents the location of the dispersal barrier. The slanted line is a hypothetical landscape with a linear gradient of precipitation. For homogeneous dispersal landscape, the savanna-forest boundary equilibrates (blue dot) at its bioclimatic limit $P_{M}$. However, in the presence of a dispersal barrier, the boundary pins near the barrier without reaching its bioclimatic limit $P_{M}$. We use $d=15$ and $D=1000$.

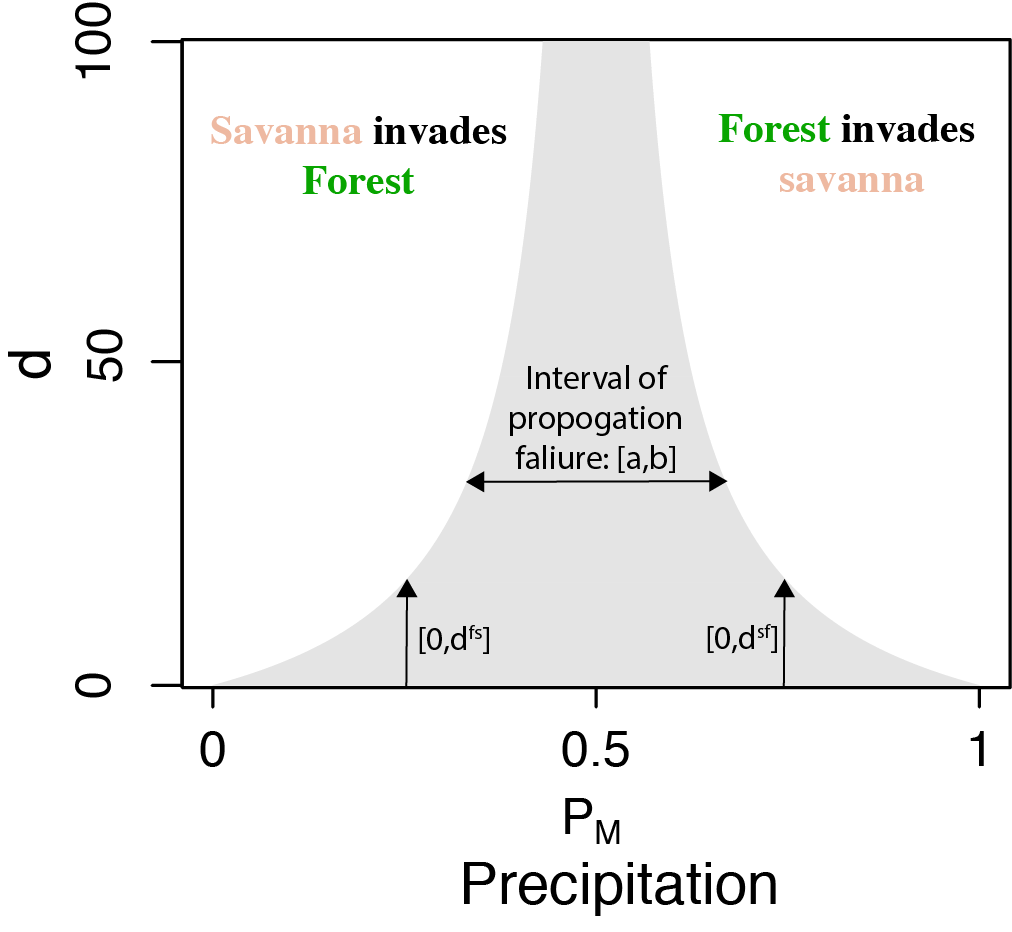

**Figure S3:** Phase plot of the direction of movement of the savanna-forest boundary at the barrier as a function of precipitation and $d$ for Eq. S1. Savanna-forest boundary stabilizes at the barrier if the $d$ and $P$ falls in the grey region. Note as $P$ approaches $P_{M}$, both $d^{sf}$ and $d^{fs}$ increases. Conversely, when $d$ is low, boundary stabilizes for a wide range of rainfall conditions, i.e., [$a,b$] interval is wide. We use $D=1000$ to generate the plot.

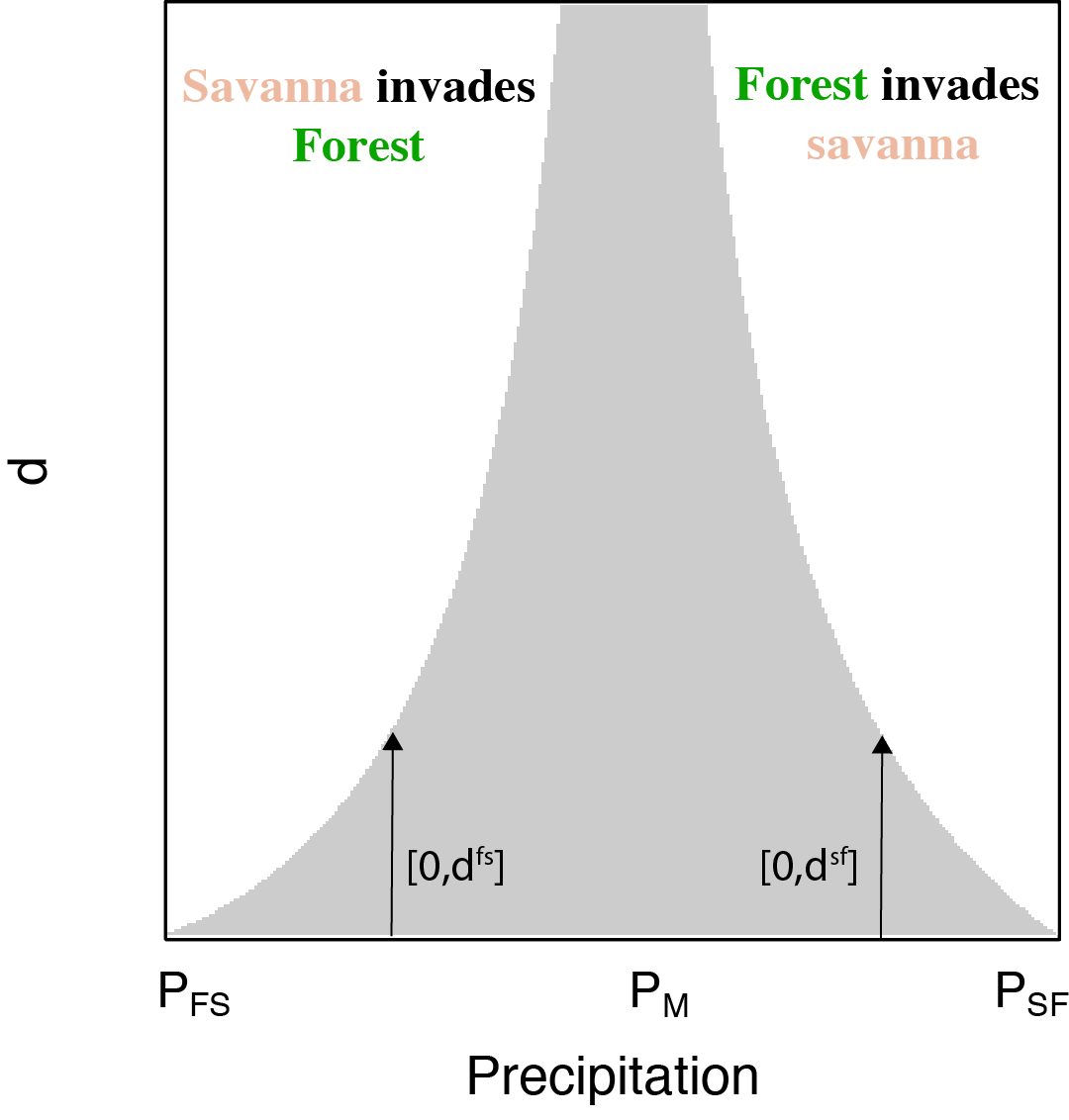

**Figure S4:** Phase plot of the direction of boundary movement at the barrier as a function of precipitation and $d$for Eq. 1 in the main text. Savanna-forest boundary stabilizes at the barrier if the $d$ and $P$ falls in the grey region. Note as $P$ approaches $P_{M}$, both $d^{sf}$ and $d^{fs}$ increases.

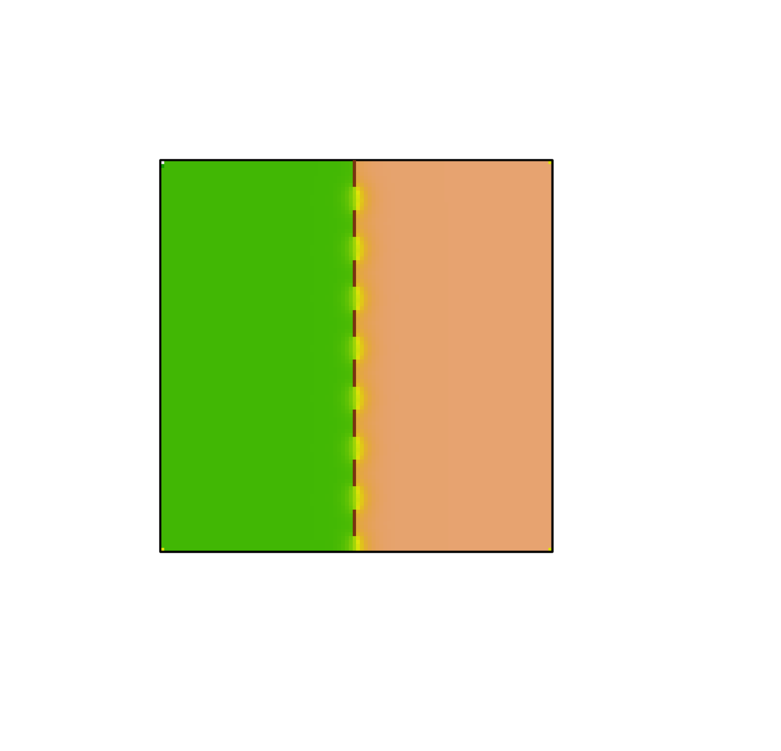

**Figure S5:** Distribution of savanna and forest in a 2D landscape with $P>P_{M}$. The brown patches show the location of limited dispersal patches. The boundary ceases to move at the barrier even though it has a small opening that could allow forest to escape. Therefore, even partial barriers can stabilize the savanna-forest boundary near the barrier.

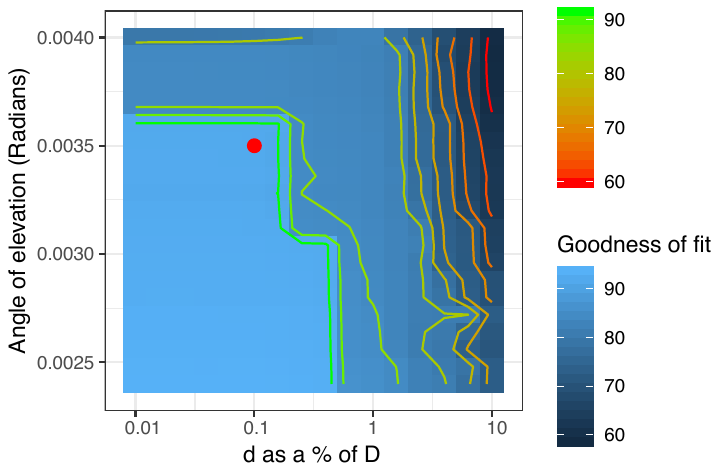

**Figure S6:** Comparing the simulated distribution of biomes with the remotely sensed biome patterns in Fig. 1A for various combinations of parameter values of $E_{th}$ and $d$. The red dot indicates the parameter values used to generate Fig 3B in the main text. The sensitivity analysis suggests that when $d$ is low (upto 1% of $D$) forest expansion into Central Madagascar might fail due to dispersal limitation. Note that $x$ axis is on a log scale.

### References

Aleman, J., B. Leys, R. Apema, I. Bentaleb, M. A. Dubois, B. Lamba, J. Lebamba, C. Martin, A. Ngomanda, L. Truc, J. M. Yangakola, C. Favier, and L. Bremond. 2012. Reconstructing savanna tree cover from pollen, phytoliths and stable carbon isotopes. Journal of Vegetation Science **23**:187-197.

Bond, W. J., J. A. Silander Jr, J. Ranaivonasy, and J. Ratsirarson. 2008. The antiquity of Madagascar’s grasslands and the rise of C4 grassy biomes. Journal of Biogeography **35**:1743--1758.

Boutton, T., S. Archer, and A. Midwood. 1999. Stable isotopes in ecosystem science: structure, function and dynamics of a subtropical savanna. Rapid Communications in Mass Spectrometry **13**:1263-1277.

Burney, D. A. 1987. Late Holocene Vegetational Change in Central Madagascar. Quaternary Research **28**:130-143.

Burney, D. A. 1993. Late Holocene Environmental-Changes in Arid Southwestern Madagascar. Quaternary Research **40**:98-106.

Burney, D. A. 1999. Rates, patterns, and processes of landscape transformation and extinction in Madagascar. Pages 145--164 Extinctions in near time. Springer.

Burney, D. A., G. S. Robinson, and L. P. Burney. 2003. Sporormiella and the late Holocene extinctions in Madagascar. Proceedings of the National Academy of Sciences **100**:10800-10805.

Burns, S. J., L. R. Godfrey, P. Faina, D. McGee, B. Hardt, L. Ranivoharimanana, and J. Randrianasy. 2016. Rapid human-induced landscape transformation in Madagascar at the end of the first millennium of the Common Era. Quaternary Science Reviews **134**:92-99.

Fáth, G. 1998. Propagation failure of traveling waves in a discrete bistable medium. Physica D: Nonlinear Phenomena **116**:176--190.

Gasse, F., E. Cortijo, J.-R. Disnar, L. Ferry, E. Gibert, C. Kissel, F. Laggoun-Défarge, E. Lallier-Verges, J.-C. Miskovsky, and B. Ratsimbazafy. 1994. A 36 ka environmental record in the southern tropics: Lake Tritrivakely (Madgascar). **318**:1513--1519.

Goel, N., V. Guttal, S. Levin, and C. Staver. 2018. Dispersal increases the resilience of tropical savanna and forest distributions. bioRxiv:476184.

Green, G. M., and R. W. Sussman. 1990. Deforestation history of the eastern rain forests of madagascar from satellite images. Science **248**:212-215.

Hogg, A. G., Q. Hua, P. G. Blackwell, M. Niu, C. E. Buck, T. P. Guilderson, T. J. Heaton, J. G. Palmer, P. J. Reimer, R. W. Reimer, C. S. M. Turney, and S. R. H. Zimmerman. 2013. Shcal13 Southern Hemisphere Calibration, 0-50,000 Years Cal Bp. Radiocarbon **55**:1889-1903.

Huffman, G. J., and D. T. Bolvin. 2013. TRMM and other data precipitation data set documentation. NASA, Greenbelt, USA **28**:1.

Humphries, A. R., B. E. Moore, and E. S. Van Vleck. 2011. Front Solutions for Bistable Differential-Difference Equations with Inhomogeneous Diffusion. SIAM Journal on Applied Mathematics **71**:1374-1400.

Jarvis, A., H. I. Reuter, A. Nelson, and E. Guevara. 2008. Hole-filled SRTM for the globe Version 4. available from the CGIAR-CSI SRTM 90m Database (<http://srtm.csi.cgiar.org>).

Koechlin, J. 1972. Flora and vegetation of Madagascar. Pages 145--190 Biogeography and ecology in Madagascar. Springer.

Matsumotot, K., and D. A. Burney. 1994. Late Holocene environments at Lake Mitsinjo, northwestern Madagascar. The Holocene **4**:16--24.

Mayaux, P., V. Gond, and E. Bartholome. 2000. A near-real time forest-cover map of Madagascar derived from SPOT-4 VEGETATION data. International Journal of Remote Sensing **21**:3139-3144.

Power, M., J. Marlon, N. Ortiz, P. Bartlein, S. Harrison, F. Mayle, A. Ballouche, R. Bradshaw, C. Carcaillet, and C. Cordova. 2008. Changes in fire regimes since the Last Glacial Maximum: an assessment based on a global synthesis and analysis of charcoal data. Climate Dynamics **30**:887-907.

Virah-Sawmy, M., L. Gillson, and K. J. Willis. 2009. How does spatial heterogeneity influence resilience to climatic changes? Ecological dynamics in southeast Madagascar. Ecological Monographs **79**:557—574.

Wright, H. T., P. Vérin, Ramilisonina, D. Burney, L. P. Burney, and K. Matsumoto. 1996. The evolution of settlement systems in the Bay of Boeny and the Mahavavy River Valley, north-western Madagascar. AZANIA: Journal of the British Institute in Eastern Africa **31**:37--73.
